# Carm1 regulates the speed of C/EBPα-induced transdifferentiation by a cofactor stealing mechanism

**DOI:** 10.1101/2022.10.03.510647

**Authors:** Guillem Torcal Garcia, Elisabeth Kowenz-Leutz, Tian V. Tian, Antonios Klonizakis, Jonathan Lerner, Luisa de Andrés-Aguayo, Clara Berenguer, Marcos Plana-Carmona, Maria Vila-Casadesús, Romain Bulteau, Mirko Francesconi, Sandra Peiró, Kenneth S. Zaret, Achim Leutz, Thomas Graf

## Abstract

Cell fate decisions are driven by lineage-restricted transcription factors but how they are regulated is incompletely understood. The C/EBPα-induced B cell to macrophage transdifferentiation (BMT) is a powerful system to address this question. Here we describe that C/EBPα with a single arginine mutation (C/EBPα^R35A^) induces a dramatically accelerated BMT in mouse and human cells. Changes in the expression of lineage-restricted genes occur as early as within 1 hour compared to 18 hours with the wild type. Mechanistically C/EBPα^R35A^ exhibits an increased affinity for PU.1, a bi-lineage transcription factor required for C/EBPα-induced BMT. The complex induces more rapid chromatin accessibility changes and an enhanced relocation (stealing) of PU.1 from B cell to myeloid gene regulatory elements. Arginine 35 is methylated by Carm1 and inhibition of the enzyme accelerates BMT, similar to the mutant. Our data suggest that the relative proportions of methylated and unmethylated C/EBPα in a bipotent progenitor can determine the velocity of cell fate choice and lineage directionality.

## INTRODUCTION

The hematopoietic system is a model of choice to understand how cells diversify into different lineages (Notta et al., 2016; Orkin and Zon, 2008). Combinations of synergistic and antagonistic transcription factors (TFs) are the main drivers of cell fate decisions, activating new gene expression programs while silencing the old ones. Their balance is an important determinant, with the most highly expressed factors becoming dominant (Graf and Enver, 2009; Okawa et al., 2018; Orkin and Zon, 2008). However, whether there are other determinants that modulate the factors’ activity and thus the velocity by which a precursor chooses alternative fates remains poorly understood.

A powerful approach to study the mechanism of cell fate decisions is TF-induced lineage conversions (Graf and Enver, 2009). C/EBPα induces the efficient transdifferentiation of B and T lineage cells into monocyte/macrophages (henceforth referred as macrophages) (Laiosa et al., 2006; Xie et al., 2004). This conversion requires the transcription factor PU.1, a key component of the regulatory networks that define lymphoid and myeloid cells(Arinobu et al., 2007; Leddin et al., 2011; Singh et al., 1999). C/EBPα contains a C-terminal basic region leucine zipper DNA-binding domain (bZip) as well as an N-terminal transactivation domain divided into distinct transactivating elements (TE-I, II and III) (Ramberger et al., 2021). During hematopoiesis it is most highly expressed in granulocyte-macrophage progenitors (GMPs) (Ohlsson et al., 2016) and its ablation blocks the formation of GMPs and granulocytes while reducing the number of monocytes (Heath et al., 2004; Ma et al., 2014; Zhang et al., 2004).

Protein post-translational modifications can alter protein structure, subcellular localization and interactome and may dynamically coordinate signaling networks (Deribe et al., 2010; Torcal Garcia and Graf, 2021). Arginine methylation is a common protein modification effected by protein arginine methyltransferases (Prmts), which can catalyze asymmetrical and symmetrical arginine dimethylation, as well as monomethylation (Wu et al., 2021). While most studies on the role of arginine methylation have focused on histones it may also affect the function of proteins involved in DNA replication (Guo et al., 2010) and differentiation (Kawabe et al., 2012; Kowenz-Leutz et al., 2010). Among the Prmts, Carm1 (Prmt4) is particularly relevant for developmental decisions such as during early embryo development, adipogenesis and muscle regeneration, as well as for cancer (Kawabe et al., 2012; Kim et al., 2010; Li et al., 2013; M. E. Torres-Padilla et al., 2007; Yadav et al., 2008).

Here we describe that the methylation of a specific arginine within the transcription activation domain of C/EBPα by the arginine methyltransferase Carm1 dampens the speed by which the factor induces transdifferentiation. Mechanistically, the unmethylated form of C/EBPα accelerates BMT induction by the enhanced relocation (‘stealing) of its partner PU.1 from B cell gene regulatory regions to myeloid regions, accompanied by an accelerated closing and opening of chromatin. Our data suggest that the two forms of C/EBPα bias the differentiation of bipotent progenitors towards alternative lineages.

## RESULTS

### Mutation of arginine 35 of C/EBPα accelerates immune cell transdifferentiation

To identify post-translational modifications that are associated with the BMT-inducing ability of C/EBPα (Figure 1A), we focused on arginines in the factor’s transactivation domain. We identified three evolutionarily conserved arginines (R12, R35, and R86) located within the N-terminus (Figure 1B) in two transactivating elements (TE-I and TE-II) required for efficient BM (Stoilova et al., 2013). First we generated a triple mutant (C/EBPα^TM^) in which these arginines were substituted by alanines (Figure 1B) and inserted it into a ß-estradiol (ß-est)-inducible retroviral vector (Xie et al., 2004), generating C/EBPα^TM^-ER-GFP. This construct was used to infect bone marrow-derived B cell precursors (henceforth called B cells) grown on feeder cells for 2 days and GFP+ B cells isolated. The infected cells were re-seeded on feeders, cultures treated with ß-est and expression of the macrophage marker Mac-1 (CD11b) and the B cell marker CD19 (Springer et al., 1979; Wang et al., 2012) monitored by FACS at various days later. Surprisingly, C/EBPα^TM^ greatly accelerated BMT, generating almost 100% macrophage-like cells (Mac-1+, CD19-) within 3 days compared to 4 to 5 days for C/EBPαWT-infected cells (Figure, 1C, S1A).

**Figure 1.**
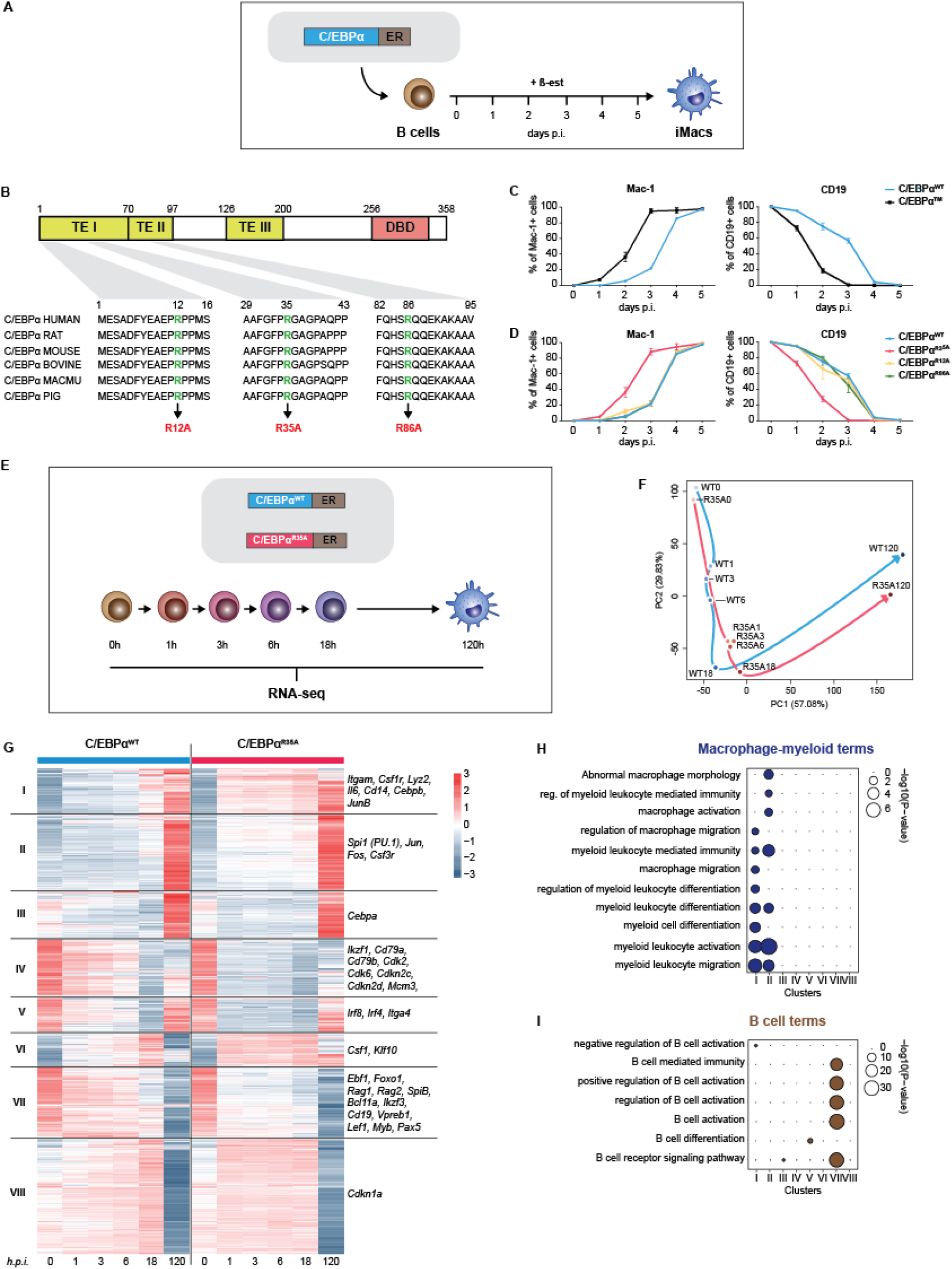
Mutation of arginine 35 in C/EBPα accelerates B cell to macrophage transdifferentiation. **A.** Schematics of the B cell to macrophage transdifferentiation (BMT) method. Bone marrow-derived pre-B cells infected with C/EBPα-ER retrovirus are treated with ß-est to induce the factor’s translocation into the nucleus, inducing a BMT within 4 to 5 days. **B.** C/EBPα structure (TE = transactivation element; DBD = DNA-binding domain) and location of conserved arginines R12, R35, and R86 within the N-terminus, which were replaced by alanines. **C.** Kinetics of BMT induced by wild type (WT) C/EBPα and a triple mutant (C/EBPα^TM^) with alanine replacements of R12, R35 and R86. BMT was assessed by Mac-1 and CD19 expression (mean ± s.d., n=3). **D**. Kinetics of BMT induced by C/EBPα^WT^ and single arginine to alanine replacements at C/EBPα R12, R35, and R86. **E.** Schematics of experimental approach for RNA-sequencing (RNA-seq) of B cells infected with either C/EBPα^WT^- or C/EBPα^R35A^- ER retroviral constructs induced for various timepoints. **F.** Principal component analysis (PCA) of 11,780 differentially expressed genes (DEGs) during BMT (n=2). Arrows connecting individual time points visualize trajectories **G.** Hierarchical clustering of DEGs with representative genes shown next to each cluster. **H-I.** Gene ontology (GO) enrichment analysis of macrophage-myeloid (**H**) and B cell (**I**) terms of the clusters from Figure 1G. Diameter of circles is proportional to the p-value. See also Figure S1.

Next, we tested the effect of alanine replacement for each of the 3 individual arginines (R12A, R35A, and R86A) and found that C/EBPα^R35A^ recapitulated the phenotype of C/EBPα^TM^, while C/EBPα^R12A^ and C/EBPα^R86A^ showed no such effect (Figures 1D, **S1A**). Five-day-induced C/EBPα^R35A^ cells resembled normal macrophages similar to those seen with C/EBPα^WT^ cells, consisting of large, mostly adherent cells, with extensive f-actin filaments and eccentric nuclei. In addition, the cells were highly phagocytic, as >90% of them ingested carboxylated beads (Figures S1B, C).

These data show that the replacement of arginine 35 with alanine in C/EBPα dramatically accelerates the factor’s capacity to induce a BMT, as evidenced by a higher velocity of silencing and activation of B cell and macrophage markers, respectively. Moreover, the induced cells resembled normal macrophages and were functional.

### C/EBPα^R35A^ hastens gene expression changes of lineage-associated genes at early time points

To study the effects of C/EBPα^R35A^ on gene expression, we performed RNA-sequencing (RNA-seq) of infected B cells induced for 0, 1, 3, 6, 18, and 120 hours (Figure 1E). Principal component analysis (PCA) showed a pronounced acceleration in the trajectory of differentially expressed genes throughout BMT (11,780 genes) compared to the WT virus. Strikingly, induction of C/EBPα^R35A^ cells for just 1 hour caused changes similar to 18 hours induced C/EBPα^WT^ cells, with their trajectories converging again at 120 hours post induction (hpi; Figure 1F). The vast majority of genes affected by the wild type and the mutant exhibited similar expression levels at the endpoint of the conversion, indicating that the mutant mostly accelerates the speed of BMT without inducing an aberrant phenotype (Figure S1D). Moreover, the largest differences in gene expression values between wild type and mutant cells were observed at 1 and 3 hpi (Figure S1D). Hierarchical clustering of all the 11,780 differentially expressed genes throughout BMT yielded 8 clusters (Figure 1G). These could be separated into two large groups, with genes in clusters I, II, IV and VIII displaying faster activation by C/EBPα^R35A^, while clusters IV, V and VII showed faster silencing. Macrophage-myeloid related GO terms were enriched in clusters I and II (Figures 1H, S1F) and included the myeloid-restricted genes *Itgam* (encoding Mac-1) *Lyz2* (lysozyme), *Csf1r* (M-CSF receptor) and *Cd14* (Figures 1G, S1G). Conversely, B cell-related GO terms were enriched in cluster VII (Figures 1I, S1F) and included the B cell-restricted genes *Cd19*, *Pax5*, *Ebf1* and *Rag2* (Figures 1G, S1G). The kinetics of individual macrophage and B cell-associated genes (Figure S1H) further illustrate the C/EBPα^R35A^-induced BMT acceleration.

These results extend the findings obtained with B cell and macrophage cell surface markers to thousands of differentially regulated lineage-associated genes. The most dramatic differences in gene expression changes induced by C/EBPα^R35A^ occurred within 3 hpi and then converged again at 120 hpi.

### C/EBPα^R35A^ accelerates chromatin remodelling at regulatory elements of lineage-restricted genes

Major gene expression changes are typically associated with extensive chromatin remodeling (Klemm et al., 2019). To study changes in chromatin accessibility occurring during BMT, we performed assays for Transposase-Accessible Chromatin using sequencing (ATAC-seq) at various time points after C/EBPα^WT^ and C/EBPα^R35A^ induction (Figure 2A).

**Figure 2.**
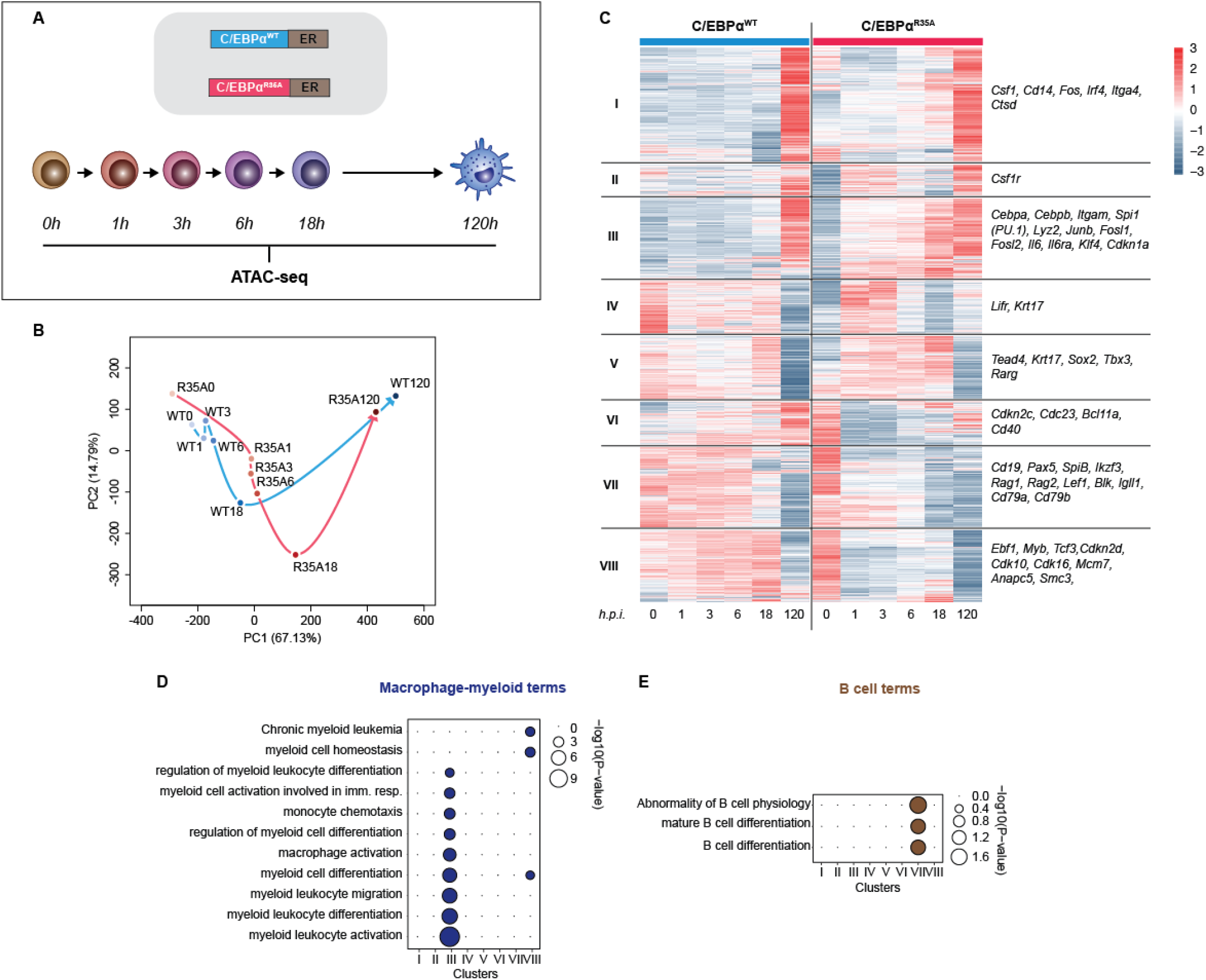
C/EBPα^R35A^ accelerates chromatin accessibility at gene regulatory elements of lineage-restricted genes. **A.** Experimental approach used for chromatin accessibility profiling. B cells infected with either C/EBPα^WT^- ER or C/EBPα^R35A^-ER retroviral constructs (n=2 biological replicates) were induced for the indicated times and processed for ATAC-seq. **B.** PCA of differential chromatin accessibility dynamics during BMT induced by C/EBPα^WT^ (cyan) or C/EBPα^R35A^ (magenta), based on 91,830 ATAC-seq peaks differentially called for the two conditions. Arrows connecting individual timepoints show trajectories. **C.** Hierarchical clustering of differentially accessible promoters (14,233 peaks) with representative genes shown next to each cluster. **D-E.** Gene ontology analysis of macrophage-myeloid (**D**) and B cell (**E**) terms of each cluster. Diameter of circles is proportional to the p-value. See also Figure S2.

ATAC-seq revealed 91,830 peaks significantly different between wild type and mutant cells in at least one time point, indicating differential chromatin accessibility. These regions fell into three groups: a) faster opening Gene Regulatory Elements (GREs), with highest peaks at 120hpi (43,429 peaks); b) faster closing GREs, with highest peaks at 0 hpi (36,380 peaks) and c) transiently opening GREs with highest peaks at 18 hpi (12,021 peaks) (Figure S2A, B). While both opening and closing GREs showed a largely accelerated trend with C/EBPα^R35A^, transiently opening GREs showed only subtle differences between the two conditions (Figure S2B). PCA analysis of the differential ATAC peaks revealed an acceleration of chromatin accessibility by C/EBPα^R35A^ (Figure 2B), with values of the 1-6 hpi C/EBPα^R35A^ samples resembling the 18 hpi C/EBPα^WT^ sample.

We then grouped the 14,233 differential peaks at promoter regions into eight clusters, with genes in clusters I, II and III exhibiting opening dynamics dramatically accelerated by C/EBPα^R35A^, while genes in clusters VII and VIII showing accelerated closing (Figure 2C). GO analysis revealed an enrichment of macrophage terms for cluster III (Figure 2D) and included the macrophage-restricted genes *Itgam* and *Lyz2* (Figure 2C). Quantification of accessibility changes in GREs (including promoters and enhancers of genes) in this cluster showed an accelerated chromatin opening by C/EBPα^R35A^ at the early timepoints (exemplified by *Itgam* and *Lyz2* in Figure S2C). Conversely, the faster closing GREs in cluster VIII were enriched for B cell-related GO terms and included the B cell genes *Cd19*, *Pax5* and *Rag2* (Figures 2C, E, S2D). Differences in chromatin accessibility at these clusters were no longer apparent at 120 hpi (Figures 2B, C).

Overall, these results indicate that C/EBPα^R35A^ is more efficient at inducing chromatin opening or closing at lineage-specific GREs compared to C/EBPα^WT^, consistent with the observed acceleration of gene expression changes (Figure 1E-I). Again, these differences are most pronounced at the earliest time points.

### Differentially opening and closing chromatin regions are enriched for PU.1 motif

To test whether the accelerated changes in chromatin accessibility are due to differential DNA binding affinities, we performed an electrophoretic mobility shift assay with both proteins, using nuclear extracts from HEK-293T cells expressing either C/EBPα^WT^ or C/EBPα^R35A^. These were incubated with an end-labeled oligonucleotide containing a palindromic C/EBPα-binding motif and run on an a native acrylamide gel. The intensity of the resulting bands corresponding to C/EBPα^WT^ and C/EBPα^R35A^ complexes were similar, indicating that the mutation does not significantly affect the DNA-binding capacity of the factor (Figure 3A).

**Figure 3.**
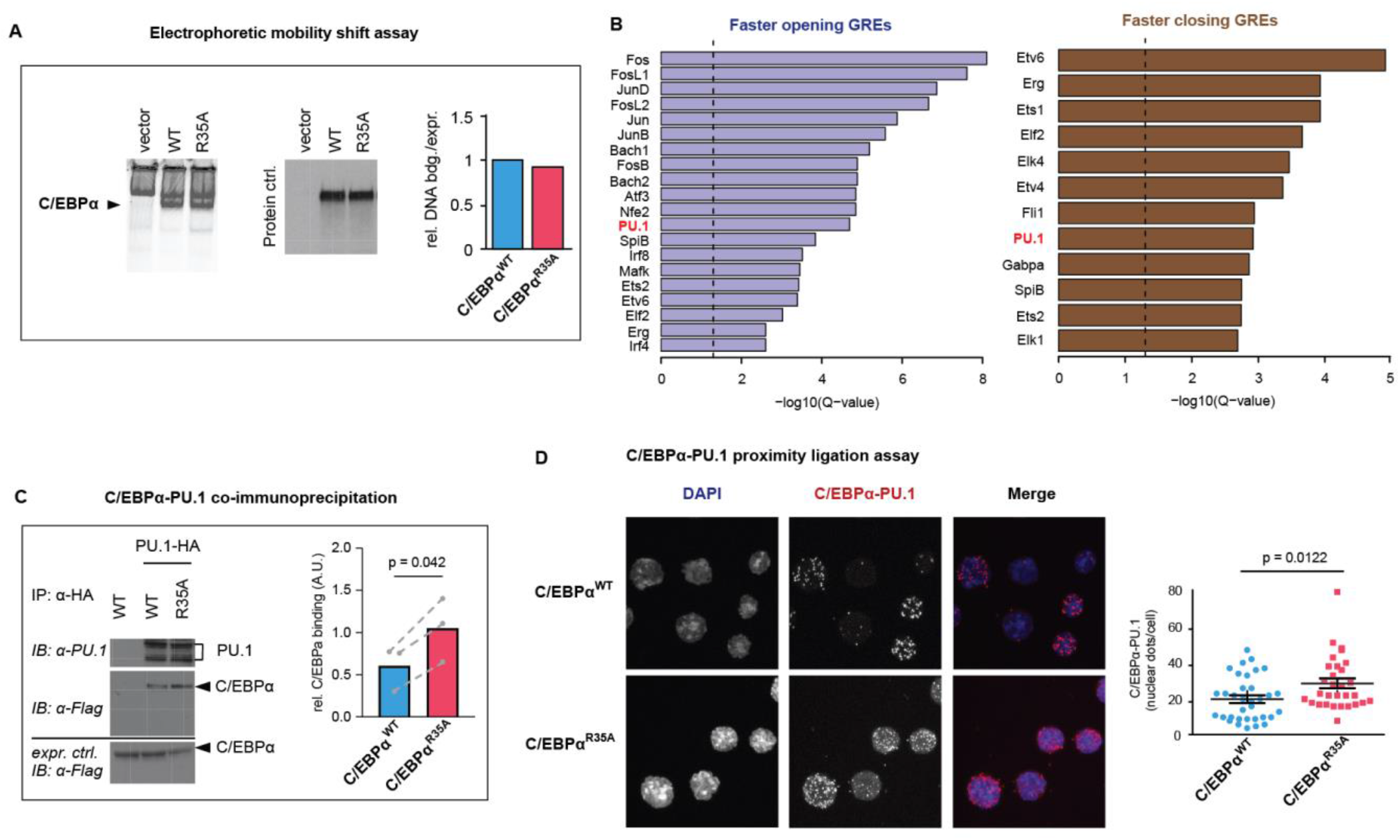
C/EBPα^R35A^ exhibits an increased affinity for PU.1. **A.** Electrophoretic mobility shift assay with nuclear extracts of HEK-293T cells transfected with either C/EBPα^WT^ or C/EBPα^R35A^ incubated with a fluorophore-labeled oligonucleotide containing a palindromic C/EBPα-binding motif (left). Protein expression control of nuclear C/EBPα proteins by western blot (middle) and densitogram-based relative DNA binding versus protein expression (right). **B.** Lists of the top *de novo* motifs in faster opening or closing GREs induced by C/EBPα^R35A^ (Figures S2A, B), with the PU.1 motif indicated in red. Dashed lines correspond to the significance threshold of Q-value (<=0.05). **C.** Co-immunoprecipitation of PU.1 and C/EBPα in HEK-293T cells transfected with either C/EBPα^WT^ or C/EBPα^R35A^ (left) and quantification of interaction of three independent experiments (right). Values shown were normalized to the expression of C/EBPα (mean + individual values). Dashed lines indicate paired values; statistical significance was determined using a paired Student’s t-test. **D.** Proximity ligation assay of C/EBPα and PU.1 in mouse B cells induced with either C/EBPα^WT^ or C/EBPα^R35A^ for 24 hours. On the left, confocal microscopy images of the cells showing nuclear dots. On the right, quantification of interactions by counting nuclear dots per cell (mean ± s.e., n=30-34; statistical significance determined using an unpaired Student’s t-test). See also Figure S3.

Another possibility is that the altered chromatin remodeling capacity of C/EBPα^R35A^ is due to the differential interaction with another protein(s). In an attempt to find such potential interactors, we performed a *de novo* motif discovery analysis with the differentially accessible GREs in the three groups by matching them against known TF motifs (Figure S2A and B). Faster and transiently opening GREs were found to be strongly enriched for AP-1/leucine zipper family TF motifs (c-Fos, c-Jun and JunB), a family of factors known to be able to heterodimerize with C/EBPα to activate myeloid genes (Cai et al., 2008). In contrast, faster closing GREs were mostly enriched for ETS family TF motifs such as Ets1, Fli1, SpiB and Gabpa, known to be associated with B cell lineage differentiation and function (Eyquem et al., 2004; Hu et al., 2001; Xue et al., 2007; Zhang et al., 2008). Several motifs were also enriched in both the accelerated chromatin opening and closing groups, including that of PU.1 and the closely related factor Spi-B (Figures 3B, S3A). Conversely, the transiently opening regions were enriched for AP-1 motifs but not for PU.1 (Figure S3B).

These observations show that chromatin regions more rapidly opened by C/EBPα^R35A^ are enriched for AP-1 family binding motifs in line with the synergism between C/EBPα and AP-1 family factors during myeloid differentiation (Cai et al., 2008). Conversely, the association of Ets family motifs with more rapidly closed regions might reflect the role of Fli1, Spi-B and in B cell differentiation (Zhang et al., 2008). That the PU.1 motif is shared between faster opening and closing regions might reflect its dual roles in the two lineages (Scott et al., 1994; Singh et al., 1999).

### C/EBPα^R35A^ exhibits an increased affinity for PU.1

Since PU.1 is a necessary partner of C/EBPα during myeloid cell specification (Heinz et al., 2010; van Oevelen et al., 2015; Xie et al., 2004) in the following we focused on the role of PU.1 during BMT. To test whether arginine 35 modulates the interaction of C/EBPα with PU.1, we performed co-immunoprecipitation experiments with cellular extracts from HEK293-T cells co-transfected with PU.1 and either WT or mutant C/EBPα. This revealed an approximately 2-fold increase in the interaction between C/EBPα^R35A^ and PU.1 compared to C/EBPα^WT^ (Figure 3C). Also, proximity ligation assays showed a stronger interaction between PU.1 and C/EBPα^R35A^ compared to C/EBPα^WT^, as determined by a significantly higher number of fluorescent nuclear dots (Figure 3D). These results therefore indicate that a mutation of C/EBPα ^R35^ increases the factor’s affinity for its obligate partner PU.1.

### C/EBPα^R35A^ shows an increased synergy with PU.1 in fibroblasts

We have previously shown that C/EBPα synergizes with PU.1 in converting NIH 3T3 fibroblasts into macrophage-like cells (Feng et al., 2008a) (Figure 4A). Therefore, to determine how the mutant behaves in this system, we generated NIH 3T3-derived cell lines (3T3aER-R and 3T3aER-A) stably expressing inducible forms of C/EBPα^WT^ or C/EBPα^R35A^, respectively. These lines were then infected with a constitutive PU.1 retroviral construct, treated with β-est, and Mac-1 levels monitored by FACS at various times post-induction. As described earlier (Feng et al., 2008b), the combination of C/EBPα^WT^ with PU.1 activated Mac-1 expression while the individual constructs did not (Figures 4B, S4A, B). Importantly, C/EBPα^R35A^ synergized with PU.1 more strongly than C/EBPα^WT^ in activating Mac-1 (Figures 4B, S4A). In addition, cells co-expressing PU.1 and C/EBPα^R35A^ exhibited dramatic morphological changes, with cells co-expressing PU.1 and C/EBPα^WT^ displaying more subtle alterations (Figure S4C).

**Figure 4.**
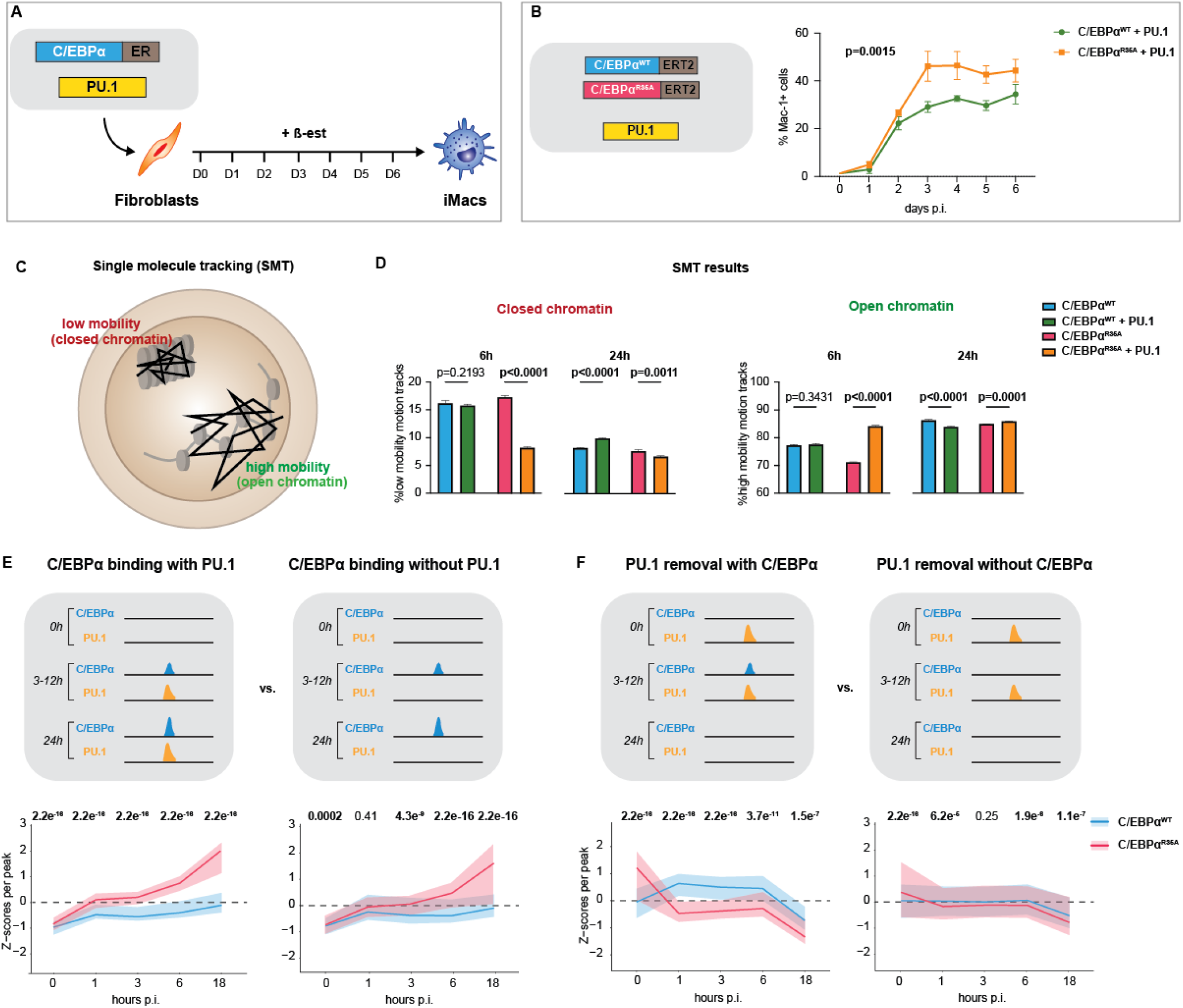
C/EBPα^R35A^ shows an enhanced synergy with PU.1 and hastens its relocation from B cell to myeloid GREs. **A.** Schematic representation of TF-induced fibroblast to macrophage transdifferentiation. NIH3T3 fibroblasts were infected C/EBPα^WT^-ER or C/EBPα^R35A^-ER in the presence or absence of PU.1 construct. Cells were induced with ß-est for the indicated times, causing a conversion to macrophage-like cells (iMacs) within 6 days p.i. **B.** Kinetics of Mac-1 expression (mean ± s.d., n=3; statistical significance was determined using two-way ANOVA). **C.** Schematic representation of single molecule movements of TFs bound to closed (low mobility) or open (high mobility) chromatin. **D.** Quantification of single cell motion tracks (mean ± s.d., n=3 x 20,000 randomized down sampled motion tracks; statistical significance determined using two-way ANOVA with multiple comparisons). **E, F.** Virtual chromatin immunoprecipitation of C/EBPα and PU.1 during BMT induced either by C/EBPα^WT^ or C/EBPα^R35A^ for the indicated times, showing schematics of peaks illustrating the different conditions tested. **E,** Selected regions corresponding to sites that are devoid of C/EBPα and PU.1 in B cells and become bound by both factors (left) or only by C/EBPα (right) throughout BMT. **F** Selected regions corresponding to sites where PU.1 is bound in B cells and either removed by transient binding of C/EBPα (left) or by another mechanism during BMT (right). See also Figure S4. Data were computed from ATAC-seq experiments (Figure 2) and from ChIP-seq of C/EBPα and PU.1 in B cells induced with ß-est for 0, 3, 12 and 24 hours (van Oevelen et al., 2015). Plots on the bottom show chromatin accessibility Z-scores per ATAC peak of B cells induced with either wild type (cyan) or mutant C/EBPα (magenta) at different hpi (line=median; shaded background=IQR; statistical significance was determined using a Wilcoxon signed-rank test).

### Single-molecule tracking experiments in fibroblasts show a PU.1-enhanced chromatin opening by C/EBPα^R35A^

To explore whether also in fibroblasts the two forms of C/EBPα exhibit differences in chromatin opening and how this is influenced by PU.1, we performed single-molecule tracking (SMT) experiments. This allows to visualize the Brownian-like movement of individual TF molecules and their interaction with open and closed chromatin (Lerner et al., 2020; Liu and Tjian, 2018) (Figure 4C). For this purpose, we generated NIH3T3 cells expressing doxycycline-inducible Halo-tagged histone H2B, C/EBPα^WT^ or C/EBPα^R35A^. After induction for either 6h or 24h these cells were used to perform SMT on ∼50 cells per condition and 20,000 single-molecule motion tracks were randomly down-sampled in triplicates to compare each condition (Chen et al., 2014). Monitoring the radius of confinement and average displacement of histone H2B allowed us to define low and high mobility chromatin, corresponding to closed and open states, respectively (Lerner et al., 2020) (Figure S4D).

Similar two-parameter assessment of motion tracks of C/EBPα^WT^ and C/EBPα^R35A^ showed that after 6 hpi, both TFs display interactions with low mobility (closed) chromatin, with C/EBPα^R35A^ showing a slightly increased interaction (Figure 4D). This observation is consistent with the elevated affinity for nucleosomes of C/EBPα measured *in vitro* (Fernandez Garcia et al., 2019; Lerner et al., 2020). At 24 hpi, both C/EBPα^WT^ and C/EBPα^R35A^ showed a decreased interaction with low mobility chromatin and increased interaction with high mobility chromatin (Figure 4D). This transition to higher mobility chromatin suggests an opening of regions bound by C/EBPα, consistent with the known pioneering function of C/EBPα (Fernandez Garcia et al., 2019).

We then tested the effect of PU.1 co-expression on interactions of C/EBPα with open or closed chromatin. At 6 hpi C/EBPα^R35A^ cells co-expressing PU.1 displayed a dramatic decrease in interaction with low mobility chromatin concomitantly with increased interaction with higher mobility chromatin, while PU.1 co-expression had little effect on the mobility of C/EBPα^WT^. This suggests a faster chromatin opening by C/EBPα^R35A^ at sites bound by PU.1 (Figure 4D and F). The observed differences between C/EBPα^WT^ and C/EBPα^R35A^ co-expressing PU.1 essentially disappeared after 24h, suggesting that the two protein complexes open closed chromatin at different speeds but reach similar endpoints (Figure 4D and E).

Altogether these results show that in 3T3 cells C/EBPα^R35A^ displays an enhanced synergism with PU.1 in that the complex induces a faster chromatin opening than the C/EBPα^WT^-PU.1 complex, coincident with stronger activation of macrophage markers and induced cell morphology changes.

### C/EBPα^R35A^ hastens the relocation of PU.1 from B cell to macrophage enhancers during BMT induction

The data described raised the possibility that C/EBPα causes a relocation of PU.1 from B cell to macrophage regulatory regions and that the mutant, through its enhanced interaction with PU.1, is more efficient at doing so. This hypothesis predicts that C/EBPα^R35A^ binding to GREs occupied by PU.1 should induce stronger changes in chromatin accessibility than C/EBPα^WT^, while sites devoid of PU.1 should behave more similarly. To test this, we performed a virtual ChIP-seq analysis of C/EBPα and PU.1 during BMT, combining previously generated ChIP-seq data (van Oevelen et al., 2015) with our new ATAC-seq data. We first identified regions stably bound by C/EBPα throughout BMT and then distinguished sites already occupied by PU.1 from PU.1-free sites. This revealed that C/EBPα^R35A^ induces a significant acceleration of chromatin *opening* at PU.1-bound regions compared to C/EBPα^WT^, while regions bound by C/EBPα alone showing much smaller differences (Figure 4E). Next, we focused on sites where PU.1 is removed by transiently bound by C/EBPα, distinguishing them from sites where PU.1 is removed yet no C/EBPα binding was detected at any timepoint. This showed that transient binding of C/EBPα^R35A^ accelerated PU.1 displacement and chromatin closing at PU.1-bound regions. In contrast, although PU.1 was also still removed at sites not targeted by C/EBPα, the effect was much milder (Figure 4F).

Altogether, our results are consistent with the hypothesis that during BMT, C/EBPα ‘steals’ endogenous PU.1 from B cell GREs and relocates it to myeloid GREs. This stealing is exacerbated by C/EBPα^R35A^, which is able to more efficiently relocate PU.1 and thus accelerate the conversion of PU.1 from a B cell regulator to a myeloid regulator, in line with the SMT results obtained in fibroblasts.

### Carm1 asymmetrically dimethylates arginine 35 of C/EBPα and decreases its affinity for PU.1

The finding that a mutation in a specific arginine of C/EBPα is responsible for the observed BMT acceleration raised the possibility that the phenotype is caused by the loss of its potential to be methylated. Since asymmetric dimethylation is one of the most common arginine modifications (Bedford and Clarke, 2009; Bedford and Richard, 2005), we first determined whether R35 is asymmetrically dimethylated. To this end, we generated two cell lines named BLaER2 and BLaER2-A, derived from the B-ALL line RCH-ACV (Jack et al., 1986) expressing the 4-hydroxytamoxifen (4-OHT)-inducible constructs C/EBPα^WT^-ERT2 and C/EBPα^R35A^-ERT2, respectively. We then induced these cells for 24h with 4-OHT, immunoprecipitated C/EBPα, and ran a Western with an antibody specific for asymmetrically dimethylated arginine (aDMA)-containing proteins. The antibody detected C/EBPα^WT^ but not C/EBPα^R35A^, thus revealing that arginine 35 is asymmetric dimethylated (Figure 5A). We next co-transfected HEK293-T cells with either C/EBPα^WT^ or C/EBPα^R35A^ and several type I Prmts, namely Prmt1, 3, 4 (Carm1) and 6; and assessed the methylation status of C/EBPα. Only Carm1 was able to induce methylation of C/EBPα^WT^ while C/EBPα^R35A^ remained unmethylated (Figures 5B, S5A).

**Figure 5.**
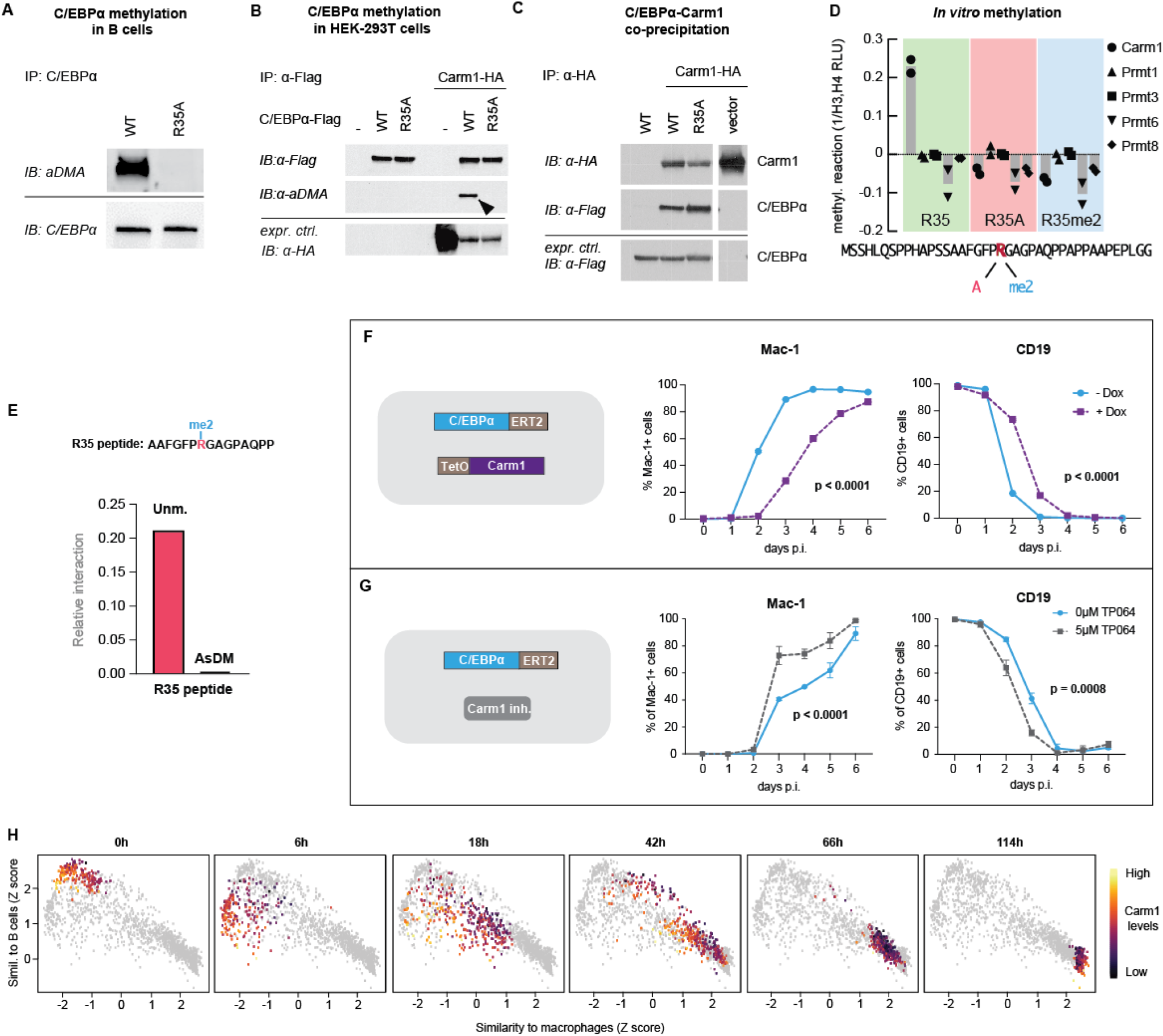
Carm1 asymmetrically dimethylates arginine 35 and regulates the speed of C/EBPα-induced BMT. **A.** Immunoprecipitation (IP) and immunoblotting (IB) of C/EBPα and asymmetrically dimethylated arginine (aDMA) containing proteins. **B.** Immunoprecipitation of C/EBPα from HEK293T cells co-transfected with either C/EBPα^WT^-Flag or C/EBPα^R35A^-Flag with or without Carm1-HA, followed by immunoblot with antibodies against aDMA, Flag and HA. **C.** Immunoprecipitation of Carm1 from HEK293T cells co-transfected with either C/EBPα^WT^-Flag or C/EBPα^R35A^-Flag and Carm1-HA, followed by immunoblot with antibodies against Flag and HA. **D.** In vitro methylation assays with recombinant Carm1, Prmt1, Prmt3, Prmt6 or Prmt8 proteins together with C/EBPα peptides (aa 15-54) that contain either unmethylated arginine 35 (green), with an alanine replacement (A, magenta), or asymmetrically dimethylated (me2, cyan) (mean and individual values are displayed, n=2). **E.** Interaction with PU.1 of a 14-mer peptide (top) containing either an unmethylated (Unm.) or an asymmetrically dimethylated arginine (me, AsDM). The data were extracted from (Ramberger et al., 2021). **F.** Effect of Carm1 overexpression on BMT kinetics of human B cells measured by Mac-1 and CD19 expression (mean ± s.d., n=3, statistical significance was determined using two-way ANOVA). **G.** Same as F, but effect of Carm1 inhibition by 5µM of TP064. **H.** Correlation of Carm1 expression levels in single cell trajectories with B cell and macrophage states. Data extracted from previously published work (Francesconi et al., 2019).

To rule out the possibility that the R35 mutation is impaired in its interaction with Carm1 we performed Co-IP experiments in HEK293-T cells co-transfected with Carm1 and either C/EBPα^WT^ or C/EBPα^R35A^, which showed that both proteins are able to interact with the enzyme (Figure 5C). To quantitatively assess the interaction of C/EBPα^WT^ and C/EBPα^R35A^ with Carm1 we performed a PLA assay. For this, NIH3T3 cell lines carrying ER fusions of C/EBPα^WT^ and C/EBPα^R35A^ were induced with β-est for 24 hours and subjected to the assay, involving staining with antibodies to C/EBPα and PU.1. We observed nuclear dots in both lines, with slightly higher numbers in C/EBPα^R35A^ cells, supporting the notion that both forms of C/EBPα can interact with Carm1 (Figure S5B).

To further assess the enzyme’s specificity, we performed an *in vitro* methylation assay using synthetic peptides (10-14-mers), covering all 20 arginine residues of C/EBPα. Only the peptide containing arginine 35 showed a methylation signal (Figure S5C). We also performed an *in vitro* methylation assay using a C/EBPα peptide spanning amino acids 15-54 and containing either unmethylated R35, asymmetrically di-methylated R35 or an alanine replacement in the presence of either Carm1, Prmt1, Prmt3, Prmt6 or Prmt8. Only Carm1 was able to methylate the peptide with the original arginine, while no methylation was detected with the other Prmts and with peptides containing methylated R35 or an alanine replacement (Figure 5D). Finally, we investigated whether the methylation status of C/EBPα affects its affinity for PU.1, analyzing the interaction data from a peptide motif-based C/EBPα interactome screen (Protein interaction Screen on Peptide Matrix, PRISMA) (Ramberger et al., 2021) comparing an unmethylated peptide with a peptide containing an asymmetrically dimethylated arginine. This showed an impaired interaction of PU.1 with the methylated compared to the unmethylated peptide (Figure 5E).

These results indicate that Carm1 selectively targets arginine 35 of C/EBPα and that the Carm1-mediated asymmetric dimethylation of this residue decreases the factor’s affinity for PU.1.

### Carm1-mediated methylation of arginine 35 modulates C/EBPα-induced BMT

To test the effect of Carm1-mediated methylation of C/EBPα on the factor’s ability to induce BMT, we performed Carm1 gain and loss of function experiments. First, we generated a stable derivative of the BLaER2 cell line (named RRC3) that contains the reverse tetracycline transactivator and a doxycycline (Dox)-inducible Carm1 construct. A Western blot confirmed robust Carm1 expression 24 hours after Dox treatment (Figure S5D). Assessing the effects of Carm1 overexpression on the kinetics of 4-OHT-induced BMT showed a dramatic delay in both Mac-1 activation and CD19 silencing (Figure 5F, S6A). Next, we tested the effect of the Carm1 inhibitor TP064 (Nakayama et al., 2018). After verifying that 5µM of the drug impairs the asymmetric dimethylation of BAF155 (Figure S5E), a known target of Carm1 (Wang et al., 2014) we found that 4-OHT-induced RRC3 cells treated with 5uM TP064 resulted in a strongly accelerated BMT (Figures 5G, S6B). In contrast, and importantly, C/EBPα^R35A^-mediated BMT was not delayed by Carm1 overexpression (Figures S5F, S6C) nor did the Carm1 inhibitor cause an acceleration (Figures S5G, S6D, E).

Our results therefore indicate that high Carm1 expression levels cause a delay in the kinetics of C/EBPα-induced BMT by methylating R35 of the wild type protein, in line with the findings obtained with C/EBPα mutant.

### Differences of endogenous Carm1 expression correlate with the speed of BMT induction

To investigate the effect of naturally occurring differences in Carm1 expression on BMT velocity, we used a previously generated single-cell gene expression dataset of cells undergoing BMT (Francesconi et al., 2019). For this, we monitored Carm1 expression during the BMT trajectory of single cells by following their similarity to either B cells or macrophages. This showed that cells with the lowest Carm1 levels were faster in acquiring a macrophage-like identity than cells with higher levels (Figure 5H). The differences leveled off between 42 and 66 hpi, suggesting that the largest differences occur in the early stages of BMT, in line with the observation that the kinetics of altered gene expression induced by C/EBPα^WT^ and C/EBPα^R35A^ differ mostly at the beginning of the process (Figure 1F). These results further support the notion that Carm1-mediated methylation of arginine 35 modulates the velocity of C/EBPα-induced BMT.

### Carm1 inhibition biases GMPs to differentiate towards macrophages

To assess the potential of Carm1 to regulate cell fate decisions during normal myelopoiesis (Figure 6A), we investigated Carm1 RNA expression levels in different myeloid precursors as well as granulocytes and macrophages, using a dataset obtained earlier (Choi et al., 2019). This revealed a gradual decrease of Carm1 during the transition from common myeloid progenitors (CMPs) over GMPs to monocytes and granulocytes (Figure S7A). Next, we monitored the levels of AsDM-BAF155 as a proxy for Carm1 activity in sorted GMPs, granulocytes and monocytes relative to total BAF155. We observed the highest relative levels of AsDM-BAF155 in GMPs and a 3.5- and 4.5-fold reduction in granulocytes and monocytes, respectively (Figure 6B, Figure S7B). These results suggest that Carm1 RNA levels and enzymatic activity decrease during myeloid differentiation, reaching their lowest levels at in monocyte/macrophages.

**Figure 6.**
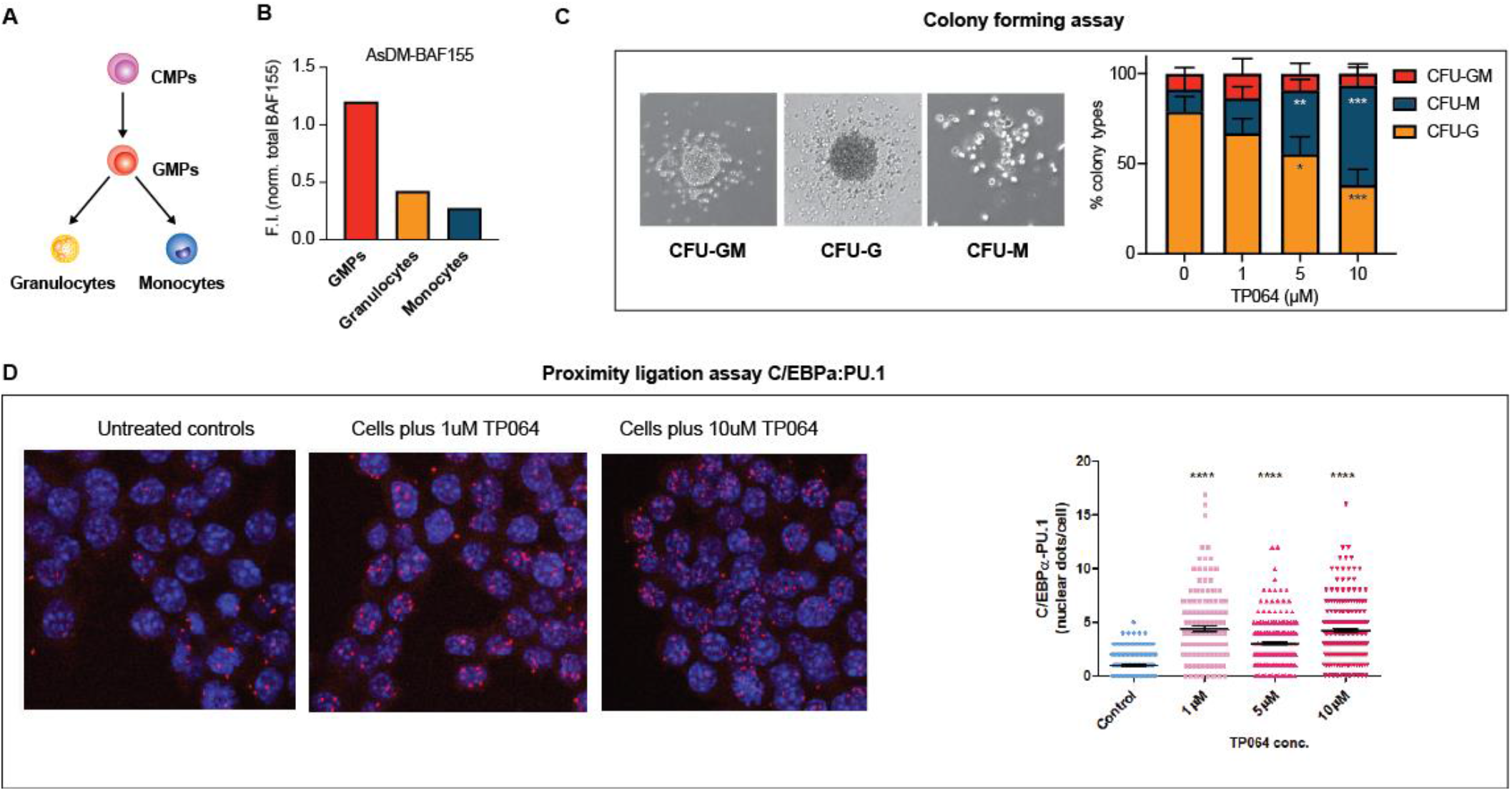
Effect of Carm1 activity on myeloid differentiation and C/EBPα-PU.1 interaction. **A.** Simplified representation of myeloid differentiation. Common myeloid progenitors (CMPs); granulocyte-macrophage progenitors (GMPs). **B.** Levels of asymmetrically dimethylated BAF155 (AsDM-BAF155) relative to total BAF155 in GMPs, granulocytes and monocytes as a proxy for Carm1 activity (see also Figure S7B). **C.** On the left, representative images of colony types obtained from GMPs grown in Methocult. On the right, quantification of colony numbers obtained in cultures without or with various concentrations of the Carm1 inhibitor TP064 for 14 days, showing percentage of bipotent (CFU-GM), monocytic (CFU-M) and granulocytic (CFU-G) colonies (mean ± s.d., n=3-4, statistical significance was determined using a one-way ANOVA for each cell type) (See also Figure S7C). **D.** Proximity ligation assay of endogenous C/EBPα and PU.1 in the mouse macrophage cell line RAW 264.7 treated for 24 hours with 1, 5 or 10uM TP064 or left untreated. On the left confocal microscopy images. On the right, counts of nuclear dots per cell (mean ± s.e., n=149-190 cells per condition). Four stars: P<0.0001 (statistical significance determined using an unpaired Student’s t-test.).

To determine whether Carm1 activity affects the decision of GMPs to differentiate into either granulocytes or monocytes, we tested the effect of the Carm1 inhibitor TP064 in a colony assay. For this, we isolated GMPs from mouse bone marrow and seeded them in a semisolid medium containing IL-3 and IL-6 in the presence of 0, 1, 2.5 or 10µM TP064. Scoring the number of the different myeloid colony types 12 days later showed a dose-dependent reduction of granulocytic colonies (CFU-G; p=0.001) and a concomitant increase of monocytic colonies (CFU-M; p=0.0003), with no effect on mixed colonies (CFU-GM; p=0.506) (Figure 6C). This bias is unlikely due to a granulocyte-selective cytotoxicity of the inhibitor since the total number of colonies remained essentially constant (Figure S7C).

Together, our results suggest that Carm1 modulates the directionality of GMPs, with unmethylated C/EBPα biasing them to differentiate towards the monocytic lineage and implying a role of methylated C/EBPα for the granulocytic lineage.

### Carm1 inhibition increases interaction between endogenous C/EBPα and PU.1

The experiments described so far, showing an increased affinity between C/EBPα with a mutated or an unmethylated R35 and PU.1, were performed after C/EBPα overexpression. To determine whether an increase in affinity can also be observed between endogenous C/EBPα and PU.1 we tested the effect of Carm1 inhibition on the mouse macrophage line RAW 264.7 (Raschke et al., 1978). For this, the cells were cultured either in the absence or in presence of 1,5 or 10uM of TP064 and subjected to a PLA assay. We observed low numbers of nuclear dots in the untreated cells and a 4 to 5 fold increase in both cultures treated with the inhibitor (Figure 6D). This increase was not due to elevated levels of the two proteins in the presence of the inhibitor, as shown by similar immunofluorescence intensities of C/EBPα and PU.1 (Figure S7D). We conclude that Carm1 inhibition increases the interaction between endogenous C/EBPα and PU.1, using a macrophage line that expresses similar levels of the two proteins.

## DISCUSSION

Here we describe a mechanism by which the speed of a hematopoietic cell fate decision is modulated. Using a model system in which C/EBPα induces a B cell to macrophage transdifferentiation (BMT) we found that an arginine 35 mutant dramatically accelerates the process. As summarized in Figure 7, our data, together with that of earlier work (van Oevelen et al., 2015) suggest that C/EBPα initiates B cell gene silencing by binding to specific GREs, a subset of which occupied by PU.1 in addition to B cell restricted regulatory factors. This binding is transient and leads to the rapid release of the complex from chromatin by an unknown mechanism. The free C/EBPα-PU.1 complex in turn translocates to macrophage-specific GREs, inducing chromatin opening and the activation of myeloid genes. During this relocation, PU.1 essentially switches from a B cell regulator to a myeloid regulator, now binding to a set of largely myeloid-specific GREs. The speed of this conversion is regulated by the levels of Carm1 in the starting cell, which determines the proportion of methylated or unmethylated arginine C/EBPα at R35. In this stealing model the C/EBPα^R35A^ mimics the unmethylated form of the factor, showing a stronger affinity for PU.1 than wild type C/EBPα. The model however, does not explain how PU.1 becomes removed from B cell GREs that are not detectably bound by C/EBPα.

**Figure 7.**
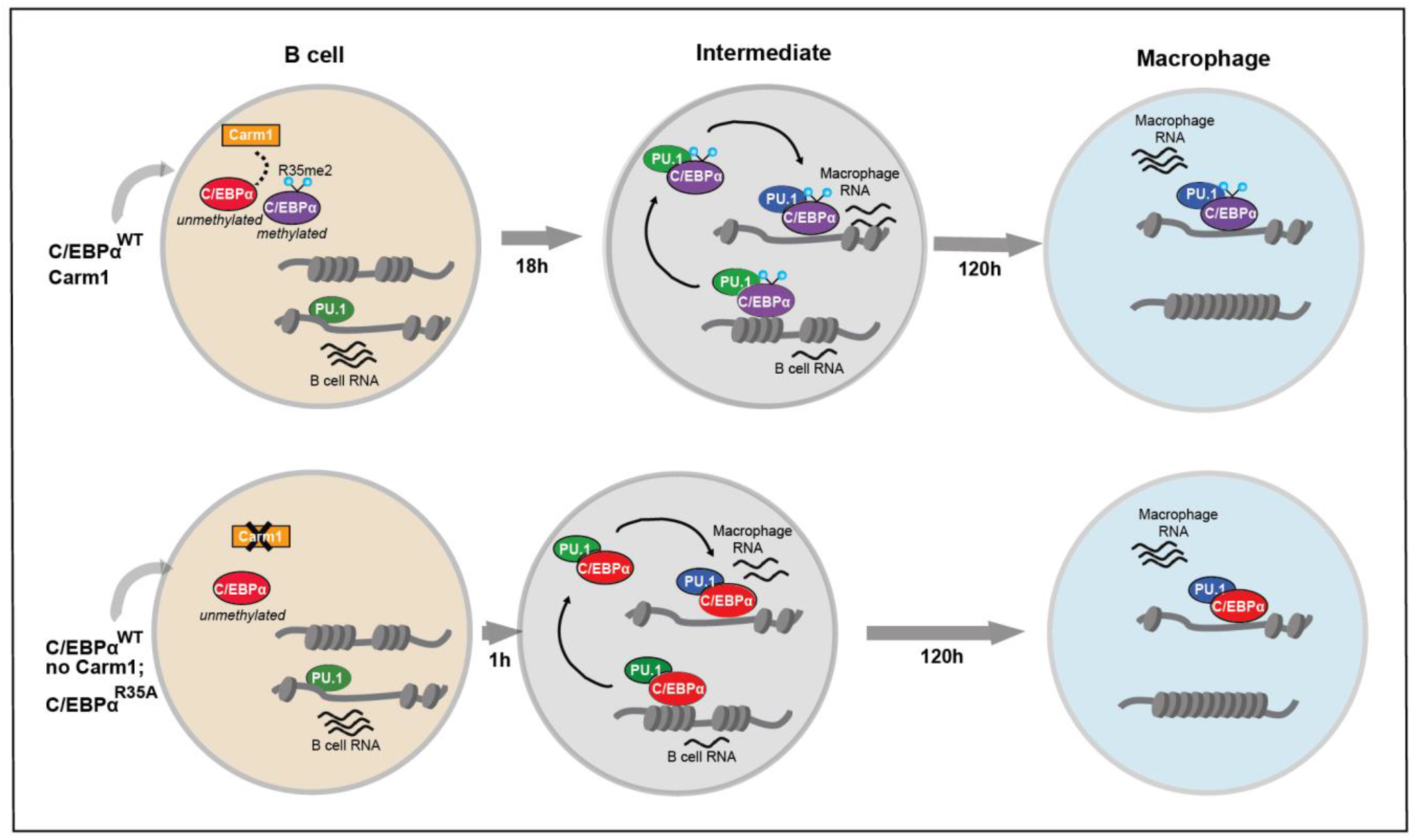
Proposed mechanism of how Carm1 modulates the velocity of BMT. The diagram shows the differences in velocity between a BMT induced by methylated and unmethylated forms of C/EBPα. In condition 1, cells are induced with C/EBPα^WT^ in the presence of Carm1 (upper part). In condition 2 (lower part) cells are induced with either C/EBPα^WT^ in the absence of Carm1 or with C/EBPα^R35A^. Note the more rapid conversion into an intermediate in the second condition. We hypothesize that C/EBPα induces gene silencing by transiently binding to gene regulatory elements (GREs) of B cells occupied by PU.1 and other B cell transcription factors. This leads to the release of the C/EBPa:PU.1 complex (and probably the other B cell factors) and chromatin closing. C/EBPα:PU.1 complexes then relocate to myeloid GREs, where they induce chromatin opening and activation of macrophage gene expression. Carm1-mediated methylation of arginine 35 delays the BMT by impairing the interaction of C/EBPα with PU.1 and relocation of PU.1 to myeloid GREs. The green symbol for PU.1 implies its role as a B cell regulator, blue as a myeloid regulator.

The observed near symmetrical acceleration of activation and silencing of B cell and myeloid-restricted genes induced by C/EBPα^R35A^ or by C/EBPα^WT^ in cells with reduced Carm1 activity suggests that PU.1 acts as a cell fate coordinator, preventing the formation of cells with aberrantly regulated lineage programs. Whether during the C/EBPα-induced BMT PU.1 acquires a different conformation when it turns from a B cell to a myeloid regulator will be interesting to determine. A critical parameter for the enhancement of myeloid differentiation during the conversion of a fetal liver T cell precursor into macrophages has been described to be cell cycle length, with cell cycle extension leading to the accumulation of high PU.1 levels (Kueh et al., 2013). Whether under physiological conditions this lengthening is induced by the activation of endogenous C/EBPα, itself known to be a potent inhibitor of the cell cycle (Nerlov, 2007), and whether it is exacerbated by a mutation of R35 remains to be studied.

A transcription factor stealing mechanism has also been described for T cell differentiation. Thus, at the DN1 progenitor stage PU.1 forms a complex with Satb1 and Runx1 at GREs of PU.1-dependent genes. Once PU.1 becomes downregulated at the DN3 stage, the associated factors are released and relocate to T cell GREs where they upregulate T cell genes (Hosokawa et al., 2018). However, in contrast to the mechanism described here, where C/EBPα acts as the ‘thief’ and PU.1 as the ‘victim’, PU.1 is the ‘thief’. In another relevant example T-bet relocates Gata3 from T_H_2 to T_H_1 genes during TH1 specification (Hertweck et al., 2022). These studies support the notion that transcription factor ‘stealing’ could be a more general mechanism by which cells coordinate silencing of the old and activation of the new differentiation program.

Remarkably, C/EBPα^R35A^ expression in B cells generates a myeloid cell-like state already within 1 hpi, only seen with the wild type after 18 hpi. Whether the observed catching up in gene expression after 120 h in C/EBPα^WT^-induced cells occurs gradually or in a more narrowly defined time window remains to be determined. Reflecting these observations, the capacity of C/EBPα to induce a transition of closed to open chromatin in fibroblasts is remarkably fast compared to other pioneer transcription factors (Lerner et al., 2020). That co-expression of PU.1 further accelerates chromatin opening in fibroblasts while activating the myeloid program suggests a powerful synergism between the two pioneer factors, regulated by methylation of arginine 35. BMT completion requires 3 to 5 days for mouse cells while human cells require 5 to 7 days (Rapino et al., 2013; Xie et al., 2004), raising the possibility that species-specific differences in Carm1 activity play a role. However, the observation that inhibition of Carm1 accelerates BMT not only in human but also in mouse cells makes this unlikely. It will be interesting to determine whether the observed species differences of BMT length reflects a higher protein stability in the human cells, as reported for neuronal specification (Rayon et al., 2020), although other mechanisms have also been described (Ebisuya and Briscoe, 2018).

In line with the results described here that Carm1 inhibition biases GMPs to differentiate into macrophage colonies, HSCs lacking Carm1 have been shown to be biased towards monocyte formation (Greenblatt et al., 2018). These observations suggest that methylated C/EBPα is required for the decision of GMPs to become granulocytes, and that this form of the factor is not simply inactivated during macrophage specification. A role of transcription factor methylation by Carm1 has also been described for muscle differentiation. Here, asymmetric dimethylation of four arginines in Pax7 enables recruitment of the MLL complex. As a consequence, Myf5 becomes transcriptionally activated, resulting in muscle cell specification (Chang et al., 2018; Kawabe et al., 2012). Carm1 has also been implicated in early embryo development and several targets have been described, including histones and chromatin modifying factors (Suresh et al., 2021; M.-E. Torres-Padilla et al., 2007), but whether this also involves the methylation of a key transcription factor is unknown.

Our observations challenge the notion that binary cell fate decisions simply result from the relative expression of antagonistic transcription factors (Graf and Enver, 2009; Moris et al., 2016). Rather, post-translational modifications, such as described here, may act as an additional regulatory layer (Torcal Garcia and Graf, 2021). Thus, the proportions of a modified versus unmodified transcription factor within a precursor population could be subject to external signaling that activates Carm1 or another enzyme that induces posttranslational modifications. Such a mechanism could operate regardless of whether binary cell fate decisions occur gradually as reported for hematopoiesis (Velten et al., 2017) or abruptly as during a neuronal differentiation cascade (Konstantinides et al., 2022). How the timing of alternative cell fate decisions is regulated is relevant not only for a better understanding of cell differentiation but also for aberrations in developmental diseases such as certain types of cancer.

## ACKNOWLEDGEMENTS

We thank the T.G. laboratory members for critical discussions, the Flow Cytometry and Microscopy units of UPF-CRG for technical assistance, the CRG Genomics core facility for sequencing and Lars Velten for feedback on the manuscript. Work in the laboratory of T. G. was supported by the Spanish Ministry of Economy, Industry and Competitiveness, (Plan Estatal PID2019-109354GB-100), the CRG, AGAUR (SGR 726) and a European Research Council Synergy grant (4D-Genome). Work in the laboratory of K.S.Z. was supported by NIH (R01GM36477).

## AUTHOR CONTRIBUTIONS

G.T.G and T.G. conceived the study and wrote the manuscript. G.T.G. performed transdifferentiation experiments (BMT and fibroblasts), cell line generation, RNA- and ATAC-seq, plasmid construction, immunofluorescence, and data analyses. E.K-L performed co-immunoprecipitation, EMSA and *in vitro* methylation assays. T.V.T. performed BMT, RNA- and ATAC-seq. A.K. and M.V-C. processed RNA-, ATAC- and ChIP-seq data. J.L. performed SMT experiments. L.A-A. performed co-immunoprecipitation, FACS, PLA and colony assays. C.B-B. performed BMT. M.P-C. confocal microscopy. R.B. and M.F. analyzed single cell expression data. S.P., K.Z., A.L. contributed ideas and discussions.

## SUPPLEMENTAL INFORMATION

**Figure S1.**
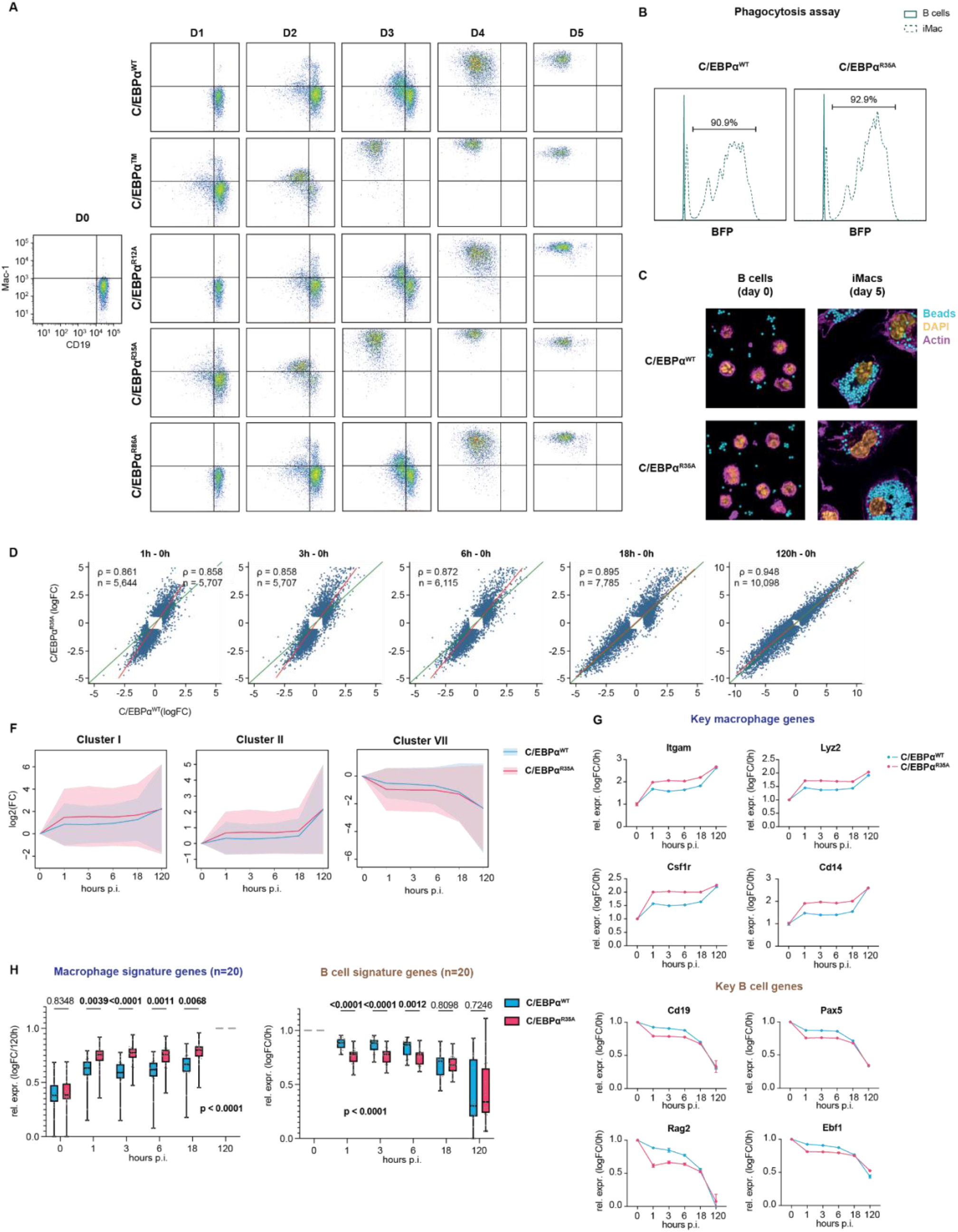
Mutation of arginine 35 in C/EBPα accelerates B cell gene silencing and macrophage gene activation. Related to Figure 1. **A.** FACS plots of Mac-1 (CD11b) and CD19 expression in B cells induced with C/EBPα^WT^, C/EBPα^TM^, C/EBPα^R12A^, C/EBPα^R35A^ or C/EBPα^R86A^ at different days p.i. **B.** Histograms showing fluorescence intensity of internalized BFP carboxylated beads in C/EBPα^WT^ and C/EBPα^R35A^-induced cells (dashed line) incubated overnight by flow cytometry. Data for uninduced control B cells are represented by a continuous line. Percentage of phagocytic cells is indicated. **C.** Immunofluorescent images of uninduced (day 0) and 5 days-induced pre B cells incubated overnight with BFP carboxylated beads. DNA was stained with picogreen (P7589) and F-actin with phalloidin Alexa Fluor 568. **D.** Scatter plots showing gene expression changes at 1, 3, 6, 18 and 120 hpi relative to 0h for B cells induced with either C/EBPα^WT^ or C/EBPα^R35A^. Red line = regression line fitted to each scatter plot; green line = identity line (x=y); ρ = Spearman correlation coefficient; n = number of differentially expressed genes. **E.** Kinetics of gene expression of clusters I, II and VII of B cells induced with either C/EBPα^WT^ (cyan) or C/EBPα^R35A^ (magenta) at different times p.i. The Y axis shows log2 fold-changes relative to uninduced cells. The lines and the shaded backgrounds correspond to the mean ± 1.64 s.d., n=1103-1868. **F.** RNA expression levels of key macrophage or B cell genes in B cells induced by either C/EBPα^WT^ (cyan) or C/EBPα^R35A^ (magenta) relative to 0h (mean ± s.d., n=2). **G.** RNA expression levels of selected macrophage and B cell signature genes in B cells induced by either C/EBPα^WT^ (cyan) or C/EBPα^R35A^ (magenta) relative to 120h and 0h, respectively (median and quartiles are represented, n=20, statistical significance was determined using multiple paired Student’s t-test for individual timepoint comparisons as well as Two-way ANOVA for overall statistical significance).

**Figure S2.**
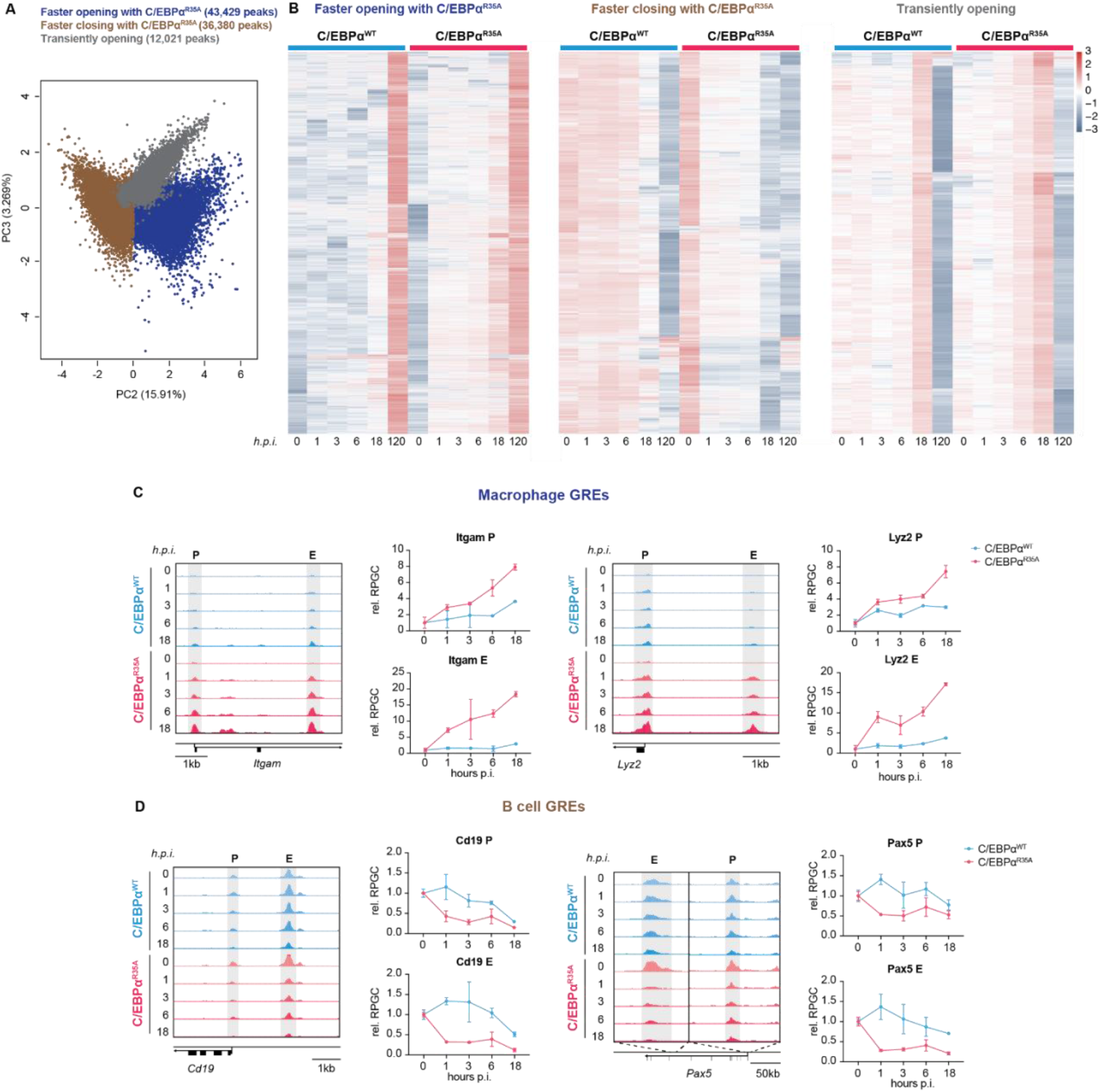
C/EBPα^R35A^ accelerates chromatin remodeling at regulatory elements of lineage-restricted genes. Related to Figure 2. **A.** PCA analysis of individual peaks showing PC2 and PC3 and the three clusters that were generated (n = 91,830 peaks). **B** Three clusters were generated from a PCA analysis shown in A. The clusters show three main trends: regions that are opened throughout BMT, more rapidly so with C/EBPα^R35A^ (blue); regions that are closed throughout BMT, also more rapidly so with C/EBPα^R35A^ (brown); and regions that peak at 18h and are closed at 120h (grey). **C.** Gene ontology (GO) enrichment of macrophage-myeloid and B cell terms of each cluster from Figure 2C. Diameter of circles is proportional to the p-value. Colored circles indicate significant enrichment. Chromatin accessibility kinetics of key macrophage (**D**) and B cell (**E**) gene regulatory elements (GREs). Genome browser views of ATAC peaks (gray highlight; P=promoter, E=enhancer) corresponding to known or putative GREs of macrophage (*Itgam and Lyz2*) and B cell genes (*Cd19 and Pax5*). Genes, direction of transcription and scale are indicated in each panel. Kinetics of chromatin accessibility at different timepoints are displayed for C/EBPα^WT^ (cyan) and C/EBPα^R35A^ (magenta) as reads per genomic content relative to 0h (RPGC).

**Figure S3.**
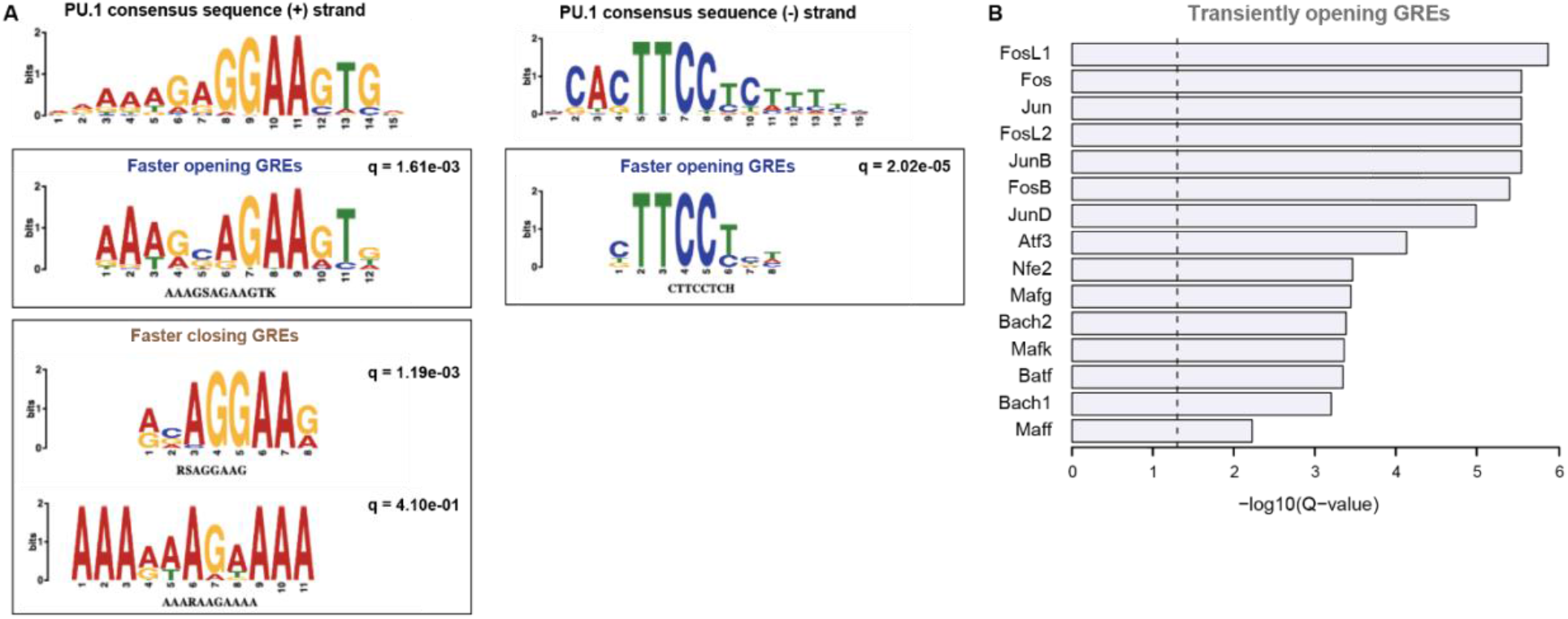
C/EBPα^R35A^ selectively interacts with PU.1. Related to Figure 3. **A.** PU.1 enriched motifs related to Figure 3D. PU.1 consensus sequence in the + and – strand is displayed (top), as well as matched enriched de novo motifs. **B.** De novo motifs matched to known TF motifs in putative in GREs that are transiently opened (grey) obtained in Figure S2A and B. Top 20 motifs are ordered by significance.

**Figure S4.**
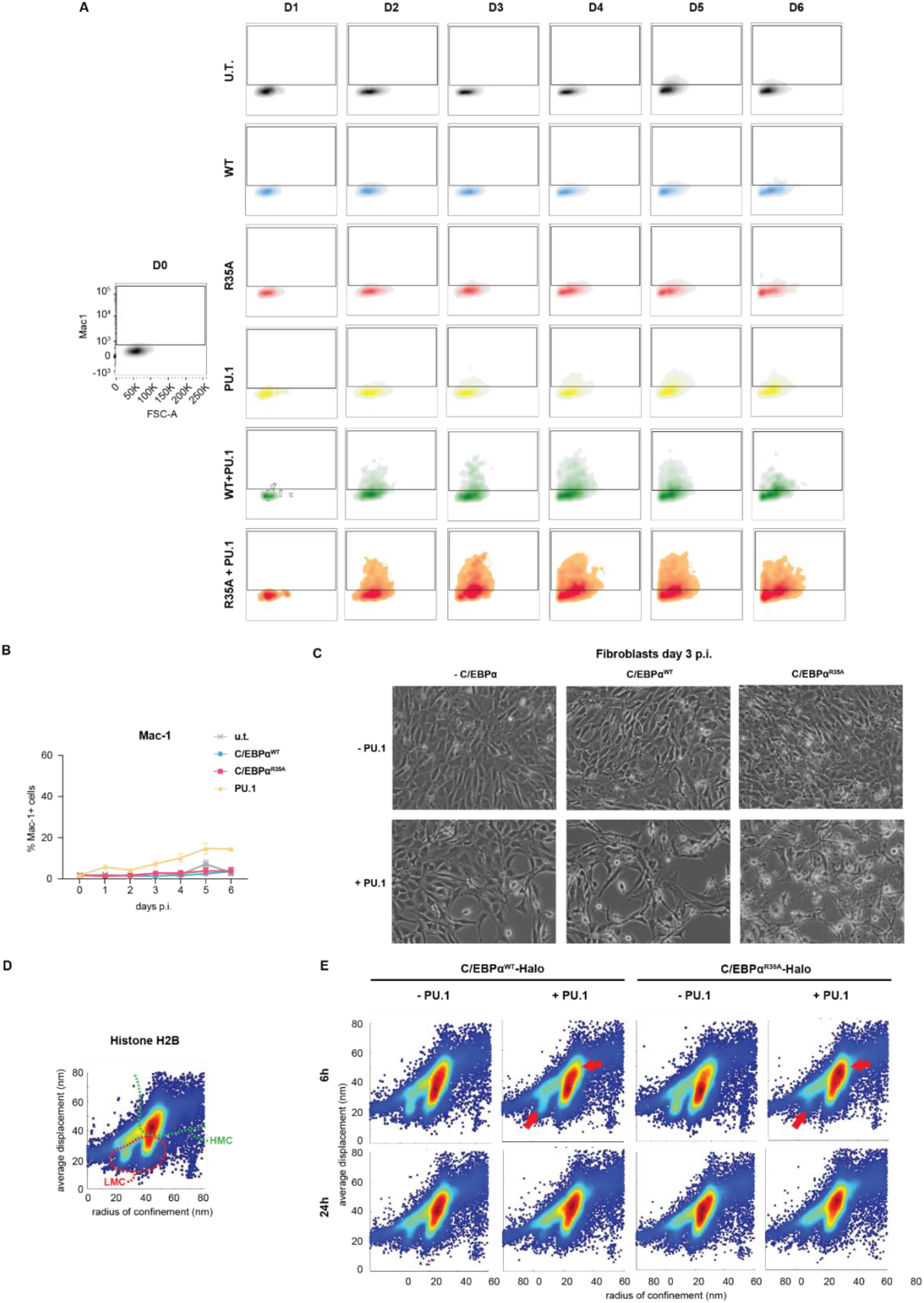
C/EBPα^R35A^ hastens the relocation of PU.1 from B cell to myeloid GREs. Related to Figure 4. **A.** FACS plots of the fibroblast to macrophage transdifferentiation by co-expression of either C/EBPα^WT^ or C/EBPα^R35A^ and PU.1 measured by Mac-1 expression by flow cytometry. **B.** Kinetics of macrophage transdifferentiation induced by C/EBPα^WT^, C/EBPα^R35A^ or PU.1 and untransduced cells (u.t.) measured by Mac-1 expression by FACS (mean ± s.d., n=3, statistical significance was determined using two-way ANOVA). **C.** Phase contrast images of NIH3T3 cells induced with either C/EBPα^WT^ or C/EBPα^R35A^ and PU.1 in different combinations for 3 days. **D.** Single molecule-tracking (SMT) of histone H2B in 3T3 cells transfected with an H2B-Halo tag construct for 24h (n = 20,000). Average displacement and radius of confinement are displayed, and chromatin mobility groups were identified (vL = very low; L = low; I = intermediate; H = high). **E.** Single molecule-tracking (SMT) of either C/EBPα^WT^ or C/EBPα^R35A^ in 3T3 cells infected with a Dox-inducible C/EBPα-Halo constructs for either 6 or 24h with or without PU.1 co-expression (n = 20,000).

**Figure S5.**
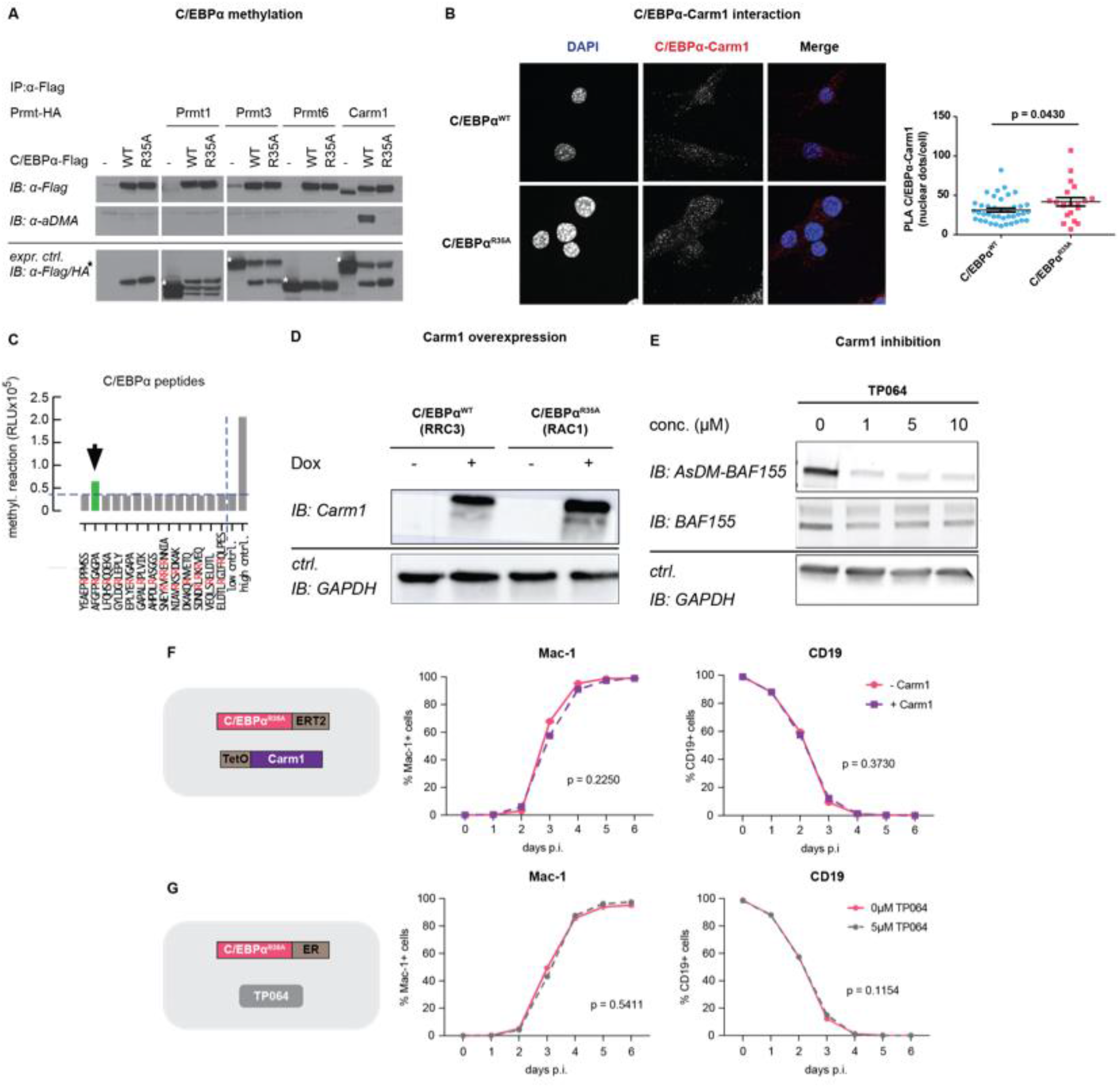
Carm1-mediated methylation of arginine 35 regulates interaction between the two proteins and the speed of C/EBPα-induced BMT. Related to Figure 5. **A.** Immunoprecipitation of C/EBPα^WT^ or C/EBPα^R35A^ (Flag) from HEK293-T cells co-transfected with either C/EBPα^WT^- or C/EBPα^R35A^-Flag and different type I Prmt enzymes (Prmt1, Prmt3, Prmt6 and Carm1), followed by immunoblot with antibodies against asymmetrically dimethylated arginine (aDMA), Flag or HA. **B.** Proximity ligation assay of Carm1 and C/EBPα^WT^ or C/EBPα^R35A^ in fibroblast lines 3T3aER-R and 3T3aER-A, respectively, induced with ß-estradiol for 24 hours. On the left, images of the cells showing DNA stained with DAPI and interaction between C/EBPα and Carm1. On the right, quantification of interaction by nuclear dots per cell (mean ± s.e.; n=20-40, statistical significance was determined using an unpaired Student’s t-test). **C.** In vitro methylation assay using Carm1 and 13 arginine-containing peptides spanning the entire C/EBPα protein (15 peptides, 20 R-residues highlighted in red). Peptide containing R35 bar indicated in green. Low control: no enzyme; high control: optimized R-methylation peptide, provided by BPS Bioscience) **D.** Western blot of Carm1 in B cell lines RRC3 and RAC1 with or without addition of Dox. **E.** Western blot of asymmetrically dimethylated BAF155 (AsDM-BAF155) and total BAF155 (BAF155) in B cells treated with different concentrations of TP064 (1-10µM). **F.** Kinetics of C/EBPα^R35A^-mediated BMT upon Carm1 overexpression by pre-treatment with Dox for 24h measured by Mac-1 and CD19 expression by flow cytometry (mean ± s.d.; n=3, statistical significance was determined using two-way ANOVA). **G.** Kinetics of C/EBPα^R35A^-mediated BMT upon Carm1 inhibition by pre-treatment with TP064 for 24h measured by Mac-1 and CD19 expression by flow cytometry (mean ± s.d.; n=3, statistical significance was determined using two-way ANOVA).

**Figure S6.**
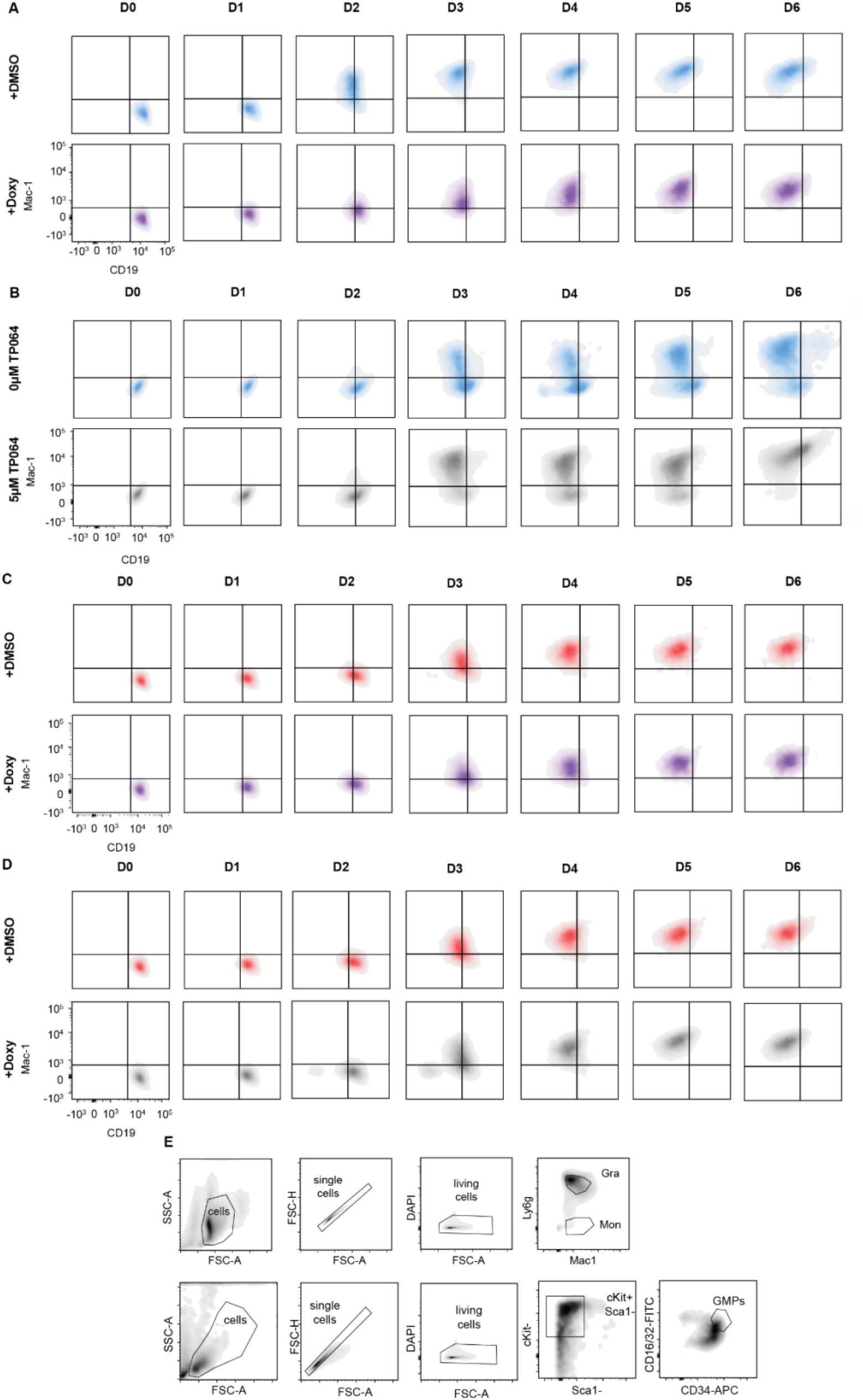
Carm1-mediated methylation of arginine 35 regulates the speed of C/EBPα-induced BMT. Related to Figure 5. **A.** FACS plots showing BMT of cells induced with C/EBPα^WT^ and exposed to Carm1 overexpression after staining for the lineage markers Mac-1 and CD19. **B.** FACS plots showing BMT of cells induced with C/EBPα^WT^ and exposed to 5µM TP064 after staining for the lineage markers Mac-1 and CD19. **C.** FACS plots showing BMT of cells induced with C/EBPα^R35A^ and exposed to Carm1 overexpression after staining for the lineage markers Mac-1 and CD19. **D** FACS plots showing BMT of induced with C/EBPα^R35A^ and exposed to 5µM TP064 after staining for the lineage markers Mac-1 and CD19. **E.** Gating strategy for sorting of bone marrow-derived granulocytes, monocytes (upper panels) and GMPs (lower panels).

**Figure S7.**
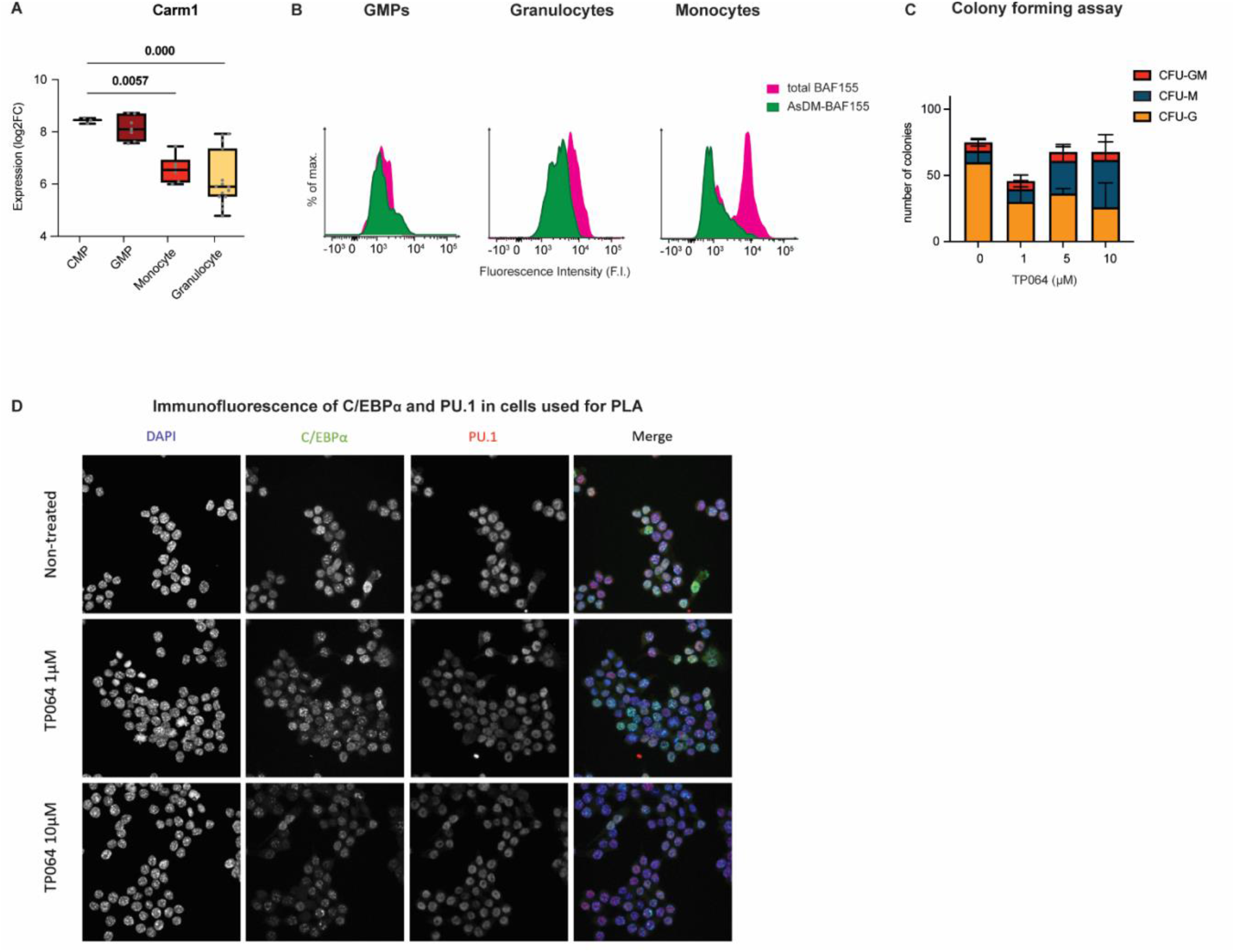
Dimethylation of C/EBPα by Carm1 is involved in the lineage choice of hematopoietic cells and C/EBPα:PU.1 interaction. Related to Figure 6. **A.** Expression of Carm1 during myeloid differentiation obtained from RNA-seq published data (Choi et al., 2019) (quartiles are represented, n=3-7, statistical significance was determined using multiple unpaired Student’s t-tests). **B.** FACS plots showing levels asymmetrically dimethylated (AsDM)-BAF155 and total BAF155 in GMPs, granulocytes and monocytes. The histograms represent fluorescence of each fraction of the protein. **C.** Colony forming unit (CFU) assay of GMPs in various concentrations of the Carm1 inhibitor TP064 after 14 days in Methocult. Total number of bipotent (CFU-GM), monocytic (CFU-M) and granulocytic (CFU-G) colonies are shown (mean ± s.d., n=3-4, statistical significance was determined using a one-way ANOVA). **D**. C/EBPα and PU.1 fluorescence in RAW cells used for PLA in Figure 6. DNA was stained with DAPI, C/EBPα with AF488 and PU:1 with AF546.

## MATERIALS AND METHODS

### Mice

As a source for the B cells used in our experiments, we used C57BL/6J mice. During experiments the number of female and male mice was balanced. Mice were housed in standard cages under 12h light-dark cycles and fed *ad libitum* with a standard chow diet. All experiments were approved by the Ethics Committee of the Barcelona Biomedical Research Park (PRBB) and performed according to Spanish and European legislation.

### Cells and cell cultures

CD19+ B cells were isolated from the bone marrow with a monoclonal antibody to CD19 (BD Biosciences, Cat#553784) using MACS sorting technology (Miltenyi Biotech) as previously described (Di Stefano, 2016). Bone marrow-derived B cells were cultured on gelatinized plates containing S17 feeder cells in RPMI culture medium (GIBCO, Cat#12633012) containing 20%-FBS (GIBCO, Cat#10270-106), 100 U/mL Penicillin-100 ng/mL Streptomycin (GIBCO, Cat#15140122), 2mM L-Glutamine (GIBCO, Cat#25030081) and 0.1mM 2-Mercaptoethanol (Invitrogen, Cat#31350010)(further addressed as **mouse B cell medium**), which was further supplemented with 10 ng/mL of IL-7 (Peprotech, Cat#217-17). HEK293-T, NIH3T3 cells (and derived) and MEFs were cultured in 10% FBS (GIBCO, Cat#10270-106) DMEM (GIBCO, Cat#12491015) medium. The final culture medium also contained 100U/mL Penicillin and 100ng/mL Streptomycin (GIBCO, Cat#15140122), 2mM L-Glutamine (GIBCO, Cat#25030081) and 0.1mM 2-Mercaptoethanol (Invitrogen, Cat#31350010) (further addressed as DMEM complete medium). RCH-ACV (and derived) human B cells were grown in RPMI culture medium (GIBCO, Cat#22400089) containing 20%-FBS (10270-106, GIBCO) (further addressed as human B cell medium).

### Induction of mouse B cell to macrophage transdifferentiation

Induction of transdifferentiation of primary pre/pro B cells (heretofore referred as B cells) isolated from the bone marrow of C57BL/6J mice was performed as previously described (Xie et al., 2004a). Briefly, B cells isolated from 8-16 weeks C57BL/6J mice were infected with C/EBPα-ER-hCD4 retrovirus, plated at 500 cells/cm^2^ in gelatinized plates (12 wells) onto mitomycin-C (Sigma, Cat#M0503)-treated MEFs (10μg/mL mitomycin-C for 3 hours to inactivate MEFs). Cells were transdifferentiated in mouse B cell medium, which was further supplemented with 10 ng/mL each of IL-7 (Peprotech, Cat#217-17), IL-3 (Peprotech, Cat#213-13), FLT-3 (Peprotech, Cat#250-31), mCSF-1 (Peprotech, Cat#315-03B), mSCF (Peprotech, Cat#250-03) and 100 nM β-estradiol (Merck Millipore, Cat#3301) to shuttle C/EBPα into the cell nucleus. Culture medium was renewed every 2 days with the same composition but without IL-7.

### Induction of fibroblast to macrophage transdifferentiation

Fibroblast transdifferentiation into macrophage experiments were performed as previously described (Feng et al., 2008b). Briefly, NIH 3T3 fibroblasts were infected with C/EBPα-ER-IRES-hCD4 retrovirus and hCD4 positive cells were sorted and a cell line was established. Cells were plated at 200,000 cells/ml in gelatinized 6-well plates and infected with PU.1ΔPEST-IRES-GFP retrovirus. After 24 hours cells were re-plated at 30,000 cells/ml in gelatinized 24-well plates in DMEM complete medium supplemented with IL-3 (Peprotech, Cat#213-13) mCSF-1 (Peprotech, Cat#315-03B) and 100 nM β-estradiol (Merck Millipore, Cat#3301) to shuttle C/EBPα into the nucleus.

### Induction of human B cell to macrophage transdifferentiation

Transdifferentiation of human B cells from the B lymphoblastic leukemia cell line RCH-ACV was performed as previously described (Rapino et al., 2013b). Briefly, RCH-ACV cells were infected with C/EBPα-ER-IRES-GFP retroviruses and GFP-positive cells were sorted, and clonal lines (BLaER2 and BLaER2-A) were generated. These lines were then infected with rtTA-Puromycin retroviruses and selected with 1µg/mL of Puromycin for 1 week. Selected cells were further infected with pHAGE-TetO-Carm1-IRES-dTomato lentiviruses. Cells were grown in human B cell medium, supplemented with 2 μg/mL of doxycycline (Sigma, Cat#D9891). Tomato-positive cells were sorted, and clonal cell lines were established (RRC3 and RAC1). For transdifferentiation cells were grown in human B cell medium, which was further supplemented with 10 ng/mL each of IL-3 (Peprotech, Cat#200-03), CSF-1 (Peprotech, Cat#315-03B) and 2.5 μM 4-hydroxytamoxifen (4-OHT) (Sigma, Cat#H7904) to shuttle C/EBPα into the cell nucleus.

### Hematopoietic colony forming assay

Bone marrow-derived GMPs from C57BL/6J mice were isolated by FACS sorting and cultured in Methocult GF M3434 (03434, Stem Cell Technologies) for 14 days. Cells were harvested from the Methocult cultures, and colonies were investigated by microscopy.

### Cell transfection

HEKT-293T cells were transfected with C/EBPα WT or mutant expression vectors in the absence or presence of PRMT1-HA, PRMR3-HA, CARM1-HA, PRMT6-HA or Pu.1 as indicated using Polyethylenimine according to the manufacturer’s protocol (PEI, Polysciences, Cat#24765-2)

### Lentivirus production and infection

Lentiviruses were produced by transfecting HEK-293T cells with 6μg of pCMV-VSV-G, 15μg of pCMVDR-8.91, and 20μg of the lentiviral vector using the calcium phosphate transfection method. Briefly, calcium phosphate-DNA precipitates were prepared by pooling the upper amounts of the three plasmids in a 2.5M CaCl2 aqueous solution. While vortexing, one volume of HBS 2X (HEPES-buffered saline solution pH=7.05, 280mM NaCl, 0.05M HEPES and 1.5mM Na2HPO4) was added dropwise to an equal volume of the calcium phosphate-DNA solution.

The mixture was incubated for 15 minutes at room temperature and added dropwise to HEK-293T cells grown in DMEM complete medium onto gelatin-coated 100mm dishes. After 8 hours of incubation at 37°C, the transfection medium was replaced with fresh medium and the supernatant collected after 24 hours. The medium was replaced again, and a second round of supernatant was collected after another 24 hours and mixed with the previous batch. The combined supernatants were centrifuged for 5 min at 300 rcf and filtered through 0.45μm strainers to remove cell debris. Lentiviral particles were then concentrated by centrifugation for 2 hours at 20,000 rcf (Beckman Coulter, Optima L-100K) in round bottom polypropylene tubes (Beckman Coulter, Cat#326823). After discarding the supernatants, the lentiviral pellets obtained from one 150mm dish were thoroughly re-suspended in 80 μL of PBS. 10^6^ fresh cells were then collected in 900μL of the respective culture medium and 10μL of lentiviral suspension were added. Subsequently, the virus-cell mixture was centrifuged at 1,000 rcf for 2 hours at 32°C (Beckman Coulter, Allegra X-30R). Infected cells were then cultured as described above and subsequently FACS-sorted for the establishment of clonal cell lines.

### Retrovirus production and infection

Retrovirus constructs were generated as described before (Bussmann et al., 2009). For production of virus for mouse cells platinum E cells (Cell Biolabs, Cat#RV-101) were transfected. Platinum A cells (Cell Biolabs, Cat#RV-102) were transfected for human cells. Infection of cells was performed as previously described (Di Stefano et al., 2014).

### Carm1 inhibition experiments with TP064

TP064 (Bio-Techne RD Systems, Bristol, UK) was used to inhibit Carm1 activity as previously described (Nakayama et al., 2018). For experiments with B cells, these were pre-incubated with 5µM of TP046 24 hours prior to induction with ß-est, and treatment with the inhibitor continued during the time of induction. For the colony forming assay with GMPs, 1-10µM of TP064 was added to the medium at the time of plating.

### Cell purification

Mouse bone marrow cell extraction was performed as previously described (Di Stefano et al., 2014). Briefly, femurs and tibias of C57BL/6J mice were extracted and crushed on a mortar in PBS supplemented with 4%FBS and 2 mM EDTA and filtered through 0.45μm strainers (Merck Millipore, Cat#SLHV033RB). For B cells, bone marrow-derived cells were incubated with sequentially 0.1µg per 1 million cells of both Fc block and Cd19-Biotin antibody for 10 and 20 minutes respectively, followed by 10 µL of magnetic streptavidin microbeads (Miltenyi, Cat#130-048-101) for an additional 20 minutes. Cd19+ cells were sorted using LS columns (Miltenyi, Cat#130-042-401). For B cell to macrophage transdifferentiation Cd19+ B cells were infected with C/EBPα-ER-IRES-hCD4 (WT and mutants) and cultured over MEF feeder cells for 4 days. Cultured B cells were incubated sequentially with 0.1µg per 1 million cells of both Fc block and hCD4-Biotin antibody for 10 and 20 minutes respectively, followed by 10 µL of magnetic streptavidin microbeads (Miltenyi, Cat#130-048-101) for an additional 20 minutes. hCD4+ cells were enriched with LS columns (Miltenyi, Cat#130-042-401).

For granulocytes and monocytes, bone marrow-derived cells were incubated sequentially with 0.1µg per 1 million cells of both Fc block and Mac1-Biotin antibody for 10 and 20 minutes respectively, followed by 10 µL of magnetic streptavidin beads (Miltenyi, Cat#130-048-101) for an additional 20 minutes. Mac1+ cells were sorted using LS columns (Miltenyi, Cat#130-042-401) and incubated with Mac1-PE and Ly6g-APC for 20 minutes. Mac1+ Ly6g-(monocytes) and Mac1+ Ly6g+ cells (granulocytes) were sorted using either FACS Aria or Influx cell sorters.

For granulocyte-monocyte progenitors (GMPs), bone marrow-derived cells were lineage-depleted using a Lineage Cell Depletion Kit (Miltenyi, Cat#130-090-858). Lineage negative cells were then incubated with Cd34-APC, cKit-APC-Cy7, Sca1-PE-Cy7 and Cd16/32-FITC for 1.5 hours. Sca1-cKit+ Cd34+ Cd16/32+ cells (GMPs) were sorted using either FACS Aria or Influx cell sorters.

For 3T3 NIH fibroblasts cells infected with C/EBPα-ER-IRES-hCD4 (WT and T35A) were incubated with 0.1µg per 1 million cells of both Fc block and hCD4-Biotin antibody (BD Pharmingen, Cat#555347) for 10 and 20 minutes respectively, followed by 10 µL of magnetic streptavidin beads (Miltenyi, Cat#130-048-101) for an additional 20 minutes. hCD4+ cells were purified using LS columns (Miltenyi, Cat#130-042-401).

For B lymphoblastic leukemia cells (RCH-ACV) cells stably infected with C/EBPα-ERT2-IRES-GFP, rtTA-Puro and TetO-Carm1-IRES-TdTomato were induced with 1µg/ml of doxycycline (Sigma-Aldrich, Cat#D9891). GFP+ and TdTomato+ cells were single cell-sorted using either FACS Aria or Influx cell sorters.

In co-cultures between B cells and feeder cells, non-adherent cells were collected, and joined with trypsinized adherent cells centrifuged at 300 RCF for 5 minutes. Cells were re-suspended in 100 μL PBS containing 1 μg/mL of mouse Fc block for 10 minutes. Conjugated primary antibodies were added to the blocking solution and cells were further incubated at 4°C in the dark for 20 minutes. Cells were washed with additional 1mL of PBS and centrifuged at 300 rcf for 5 minutes. The supernatant was discarded and cells were re-suspended in 500 μL of PBS containing 5 μg/mL of DAPI. Samples were processed in a FACS analyzer (LSR II, BD; Fortessa, BD) with DiVa software and data analyzed using FlowJo software.

Antibodies used for cell sorting and flow cytometry are listed in Table S1.

### Phagocytosis assay

After B cell to macrophage transdifferentiation, cells were removed from feeder cells through differential adherence to tissue culture dishes for 40 minutes. Around 200,000 of the resulting B cells (or induced macrophages) were plated in each well of a 24-well plate containing 0.01% poly-L-lysine-treated coverslips (Corning, Cat#354085) in 10% FBS-DMEM supplemented with IL-3 (Peprotech, Cat#213-13), mCSF-1 (Peprotech, Cat#315-03B) and cultured at 37°C overnight in the presence of 1:1000 diluted blue fluorescent carboxylated microspheres (Fluoresbrite, Cat#17458-10). Cells were centrifuged at 1000 RCF for 5 minutes to improve attachment to the coverslips. The supernatant was removed and the cells were washed once with PBS.

For fixation, 4% PFA was added to the wells for 20 minutes, cells were washed twice with PBS and cell membranes permeabilized with 0.1% Triton X-100 PBS (0.1% PBST) for 15 minutes at room temperature. Cells were blocked using 0.1% PBST with 3% Bovine Serum Albumin (BSA) for 30-45 minutes. Cells were washed twice in PBS. Actin filaments were subsequently stained with 1:100 diluted red phalloidin (Alexa Fluor 568, Thermo Fisher Scientific, Cat#A12380) while DNA was stained with a 1:500 diluted yellow probe (Quant-iT PicoGreen dsDNA Assay Kit, Thermo Fischer Scientific, Cat#P7589). Cells were incubated with the two dyes in 0.1% PBST containing 1% BSA at room temperature for 1 hour in the dark and washed twice with PBS afterwards. Coverslips carrying the attached cells in the well were then recovered with tweezers and mounted upside-down onto a charged glass slide containing a 14 μL drop of mounting medium (7μL Dako + 7μL 0.1% PBST). Coverslips were sealed with nail polish and imaged in a Leica TCS SPE inverted confocal microscope.

Antibodies used for immunofluorescence and intracellular staining for flow cytometry are listed in Table S2.

### Proximity ligation assay (PLA)

Proximity ligation assay was performed using Duolink Orange Kit (Sigma-Aldrich, Cat#DUO92007). Briefly, after sorting or culturing desired cell populations, 8.000 – 100.000 cells per well were seeded into 24-well plates containing 0.01% poly-L-lysine (Sigma) treated coverslips in appropriate medium, centrifuged at 1000 x g for 5 minutes and fixed with 4% PFA for 15 minutes. Subsequent steps were performed according to the kit’s protocol with antibody concentrations identical to those used for immunofluorescence. Coverslips were mounted using Fluoroshield mounting medium with DAPI (Abcam, Cat#ab104139) and imaged in a Leica TCS SPE confocal microscope.

### Intracellular staining for flow cytometry

After antibody staining of cell surface markers, cells were fixed in 4% BSA for 10 minutes at room temperature in a rotating wheel. Fixation was stopped with two washes in PBS. Cells were permeabilized in 0.1% PBST at room temperature in a rotating wheel for 10 minutes. Cells were blocked using 0.1% PBST with 3% Bovine Serum Albumin (BSA) for 30-45 minutes. Cells were washed twice in PBS. Cells were incubated with primary antibodies and secondary antibodies diluted at the stated concentrations in 0.1% PBST with 1% BSA for 2 and 1 hours, respectively, with two washes in PBS in between and after. Cells were resuspended in PBS and processed in a FACS analyzer (LSR II, BD; Fortessa, BD) with DiVa software and data analyzed using FlowJo software.

Antibodies used for immunofluorescence and intracellular staining for flow cytometry are listed in Table S2.

### Protein extraction, immunoprecipitation and Western blotting

Preparation of whole cell lysates and immunoprecipitation of WT or mutant C/EBPα proteins were performed as previously described (Kowenz-Leutz et al., 2010). Briefly, cells were lyzed (20 mM HEPES pH 7.8, 150 mM NaCl, 1 mM EDTA pH 8, 10 mM MgCl_2_, 0,1% Triton X-100, 10% Glycerol, protease inhibitor cocktail (Merck), 1mM DTT, 1mM PEFA bloc (Böhringer). Immunoprecipitation was performed with appropriate antibodies as indicated for 2 h at 4°C. Immunoprecipitated proteins were collected on Protein A Dynabeads (Invitrogen, Cat#100001D) or Protein-G Dynabeads (Invitrogen, Cat#10004D), separated by SDS-PAGE (Mini PROTEAN TGX, 4-15%, Bio-RAD #5671084). For Western blotting, samples were loaded in 10% Mini-PROTEAN TGX gels (Bio-Rad) and resolved by electrophoresis in running buffer (Table S3). Protein samples were transferred to a methanol pre-activated PVDF membrane (Bio-Rad, Cat#1620177, Bio-Rad) by running them in transfer buffer (TBS) (Table S3) for 1 hour at 300mA and 4°C. Membranes were rinsed in milliQ water and protein transfer was checked by Ponceau staining (Sigma). Transferred membranes were washed once with TBS and three times with TBS-Tween (TBST) (Table S3) followed by 5% milk in TBST for 45 min. Membranes were then incubated with primary antibodies (Table S4) in 5% milk TBST, rotating overnight at 4°C, then washed three times with TBST followed by incubation with the secondary antibodies conjugated to horseradish peroxidase in 5% milk TBST for 1 hour. After three TBST washes, proteins were detected using enhanced chemiluminescence reagents (Amersham ECL Prime Western Blotting detection) in an Amersham Imager 600 analyzer or visualized by ECL (GE Healthcare, UK).. Quantification of band intensity from scanned blots was performed with Fiji software.

### Electrophoretic mobility shift assay

Nuclear extracts were prepared from transfected HEKT cells by a mininuclear extract protocol (Schreiber et al., 1989). Electrophoretic mobility shift assays (EMSA) was performed as previously described (Kowenz-Leutz et al., 1994) using double stranded IRDye Oligonucleotides containing a C/EBP-binding site: IRD800-GACACTGGATTGCGCAATAGGCTC and IRD800-GAGCCTATTGCGCAATCCAGTGTC (Metabion). Briefly, binding reactions with nuclear extracts (2,5µg) and double stranded IRD800 oligos (20pmol) were incubated for 15 min on ice, orange loading dye (Li-Cor, Cat# P/N 927-10100) was added and protein-DNA complexes were separated on a 5% native polyacrylamide gel in 0,5x TBE at 25mA at room temperature. EMSA results were visualized and quantified (Odyssey scanner, Licor, channel 800nm).

### *In vitro* protein methylation assay

Methylation of peptides (PSL, Heidelberg, Germany, Table S5) was performed using the bioluminescence-based MTase-Glo^TM^ Assay (Promega, Cat#V7601) according to the manufacturer’s protocol. Assay conditions: 200 ng of enzyme was incubated with 5μM Peptide, 10 μM S-adenosyl-L-(methyl)-methionine as methyl donor (SAM) and 6x Methyltransferse-Glo reagent at 23^0^C for 60 minutes. S-adenosylhomocystein (SAH) generated during the reaction was converted to ADP as a proportional reaction product dependent of substrate methylation by the enzymes. Subsequent incubation with the Methyltransferase-Glo Detection Solution at 23^0^C for 30 minutes converts ADP to ATP that is used in a luciferase/luciferin-based reaction and determined as relative light units (RLU) in a Berthold luminometer (Hsiao et al., 2016).

### RNA sequencing

RNA was extracted with a miRNeasy mini kit (217004, Qiagen), quantified with a NanoDrop spectrophotometer and its quality examined in a fragment Bioanalyzer (Aligent 2100 Bioanalyzer DNA 7500 assay). cDNA was synthesized with a High-Capacity RNA-to-cDNA kit (4387406, Applied Biosystems). For RNA-sequencing (RNA-seq), libraries were prepared with a TruSeq Stranded mRNA Library Preparation Kit (Illumina) followed by single-end sequencing (50 bp) on a HiSeq2500 instrument (Illumina), obtaining at least 40 million reads per sample.

Quality control of FASTQ reads was performed using FastQC version v.0.11.3. Reads were mapped aligned to the mm10 genome using STAR version 2.5.0a (Dobin et al., 2013). Gene counts were quantified Gene expression was quantified using STAR (--quantMode GeneCounts). Normalized counts and differential gene expression analysis was carried out using DESeq2 version 1.14.1 (Love et al., 2014). For each transdifferentiation experiment, timepoint 0h was set as a reference point and any gene that exhibited a statistically significant change in expression (log2FC ≥ 0.5849625 and p-value ≤ 0.05) at a later timepoint was isolated. For PCA, log2 DESeq2 normalized counts of differentially expressed genes averaged across replicates were used. The R prcomp() command with scale=T was used. Pheatmap version 1.0.12 was used to visualize changes in gene expression for all the isolated differentially expressed genes with the following clustering options: clustering_distance_rows=”correlation”, clustering_method=”ward.D2”, scale=”row”.

#### Scatter plots

Differentially expressed genes (DEGs) were determined for each timepoint as described in the “Materials and Methods”. The union of identified DEGs in the WT and R35A systems per timepoint were used to generate scatterplots depicting the log2FC changes of the aforementioned genes for each transdifferentiation system. A regression line, colored in red, was fit for each scatterplot using the geom_smooth(method=lm) R command. The identity line (y=x line) is depicted in green. The spearman correlation coefficient (cor(method=”spearman”) function in R) and the number of DEGs are also depicted per scatterplot.

### Gene ontology analysis

Functional analyses by GO were performed with the R package “g:profiler2” version 0.2.0 (Raudvere et al., 2019). Baloonplots depict all pathways associated with a specific keyword that were found enriched in at least 1 cluster. Metaplots for each cluster depict the average log2FC values of genes per timepoint and per cluster. Shaded background corresponds to the mean values ± 1.644854 standard deviation. Gene expression analysis of signature genes was performed using the individual values of genes listed in Table S6 and normalized to timepoints 0h for B cell genes and 120h for macrophage genes.

### Chromatin accessibility by ATAC-seq

ATAC-seq was performed as published (Buenrostro et al., 2015). Briefly, cells were harvested at the mentioned timepoints, feeder-depleted and lysed and 50.000 cells used per condition. Immediately, transposition was performed using Nextera Tn5 Transposase (15027865, Illumina) at 37°C for 30 minutes. Chromatin was then purified using Qiagen MinElute PCR Purification Kit (28004, Qiagen). DNA was then amplified using NEBNext High Fidelity PCR Master Mix (M0541S, New England Biolabs Inc.) and barcoded primers (see table MMX). qPCR was performed to determine the optimal number of cycles for each condition to stop amplification prior to saturation. Quality was analyzed by gel electrophoresis and in a fragment Bioanalyzer (Agilent 2100 Bioanalyzer DNA 7500 assay).

Read quality was assessed with FastQC version v.0.11.3. Adaptors were removed using Cutadapt (version 0.4.2_dev) TrimGalore! In paired end mode (--paired –nextera)(Martin, 2011).Reads were aligned to the mm10 genome using bowtie2 (v 2.2.4) in paired end mode with standard parameters. Output SAM files were converted to BAM files using samtools (v 0.1.19) (Li et al., 2009).BAM files were sorted and indexed using the samtools commands sort and index, respectively. Low quality reads and reads associated with a not primary or supplementary alignment SAM flag were filtered out using the samtools command “samtools view -F 2304 -b -q 10”. PCR duplicates were removed with Picard MarkDuplicates (version 2.3.0) with the following options: “REMOVE_DUPLICATES=true ASSUME_SORTED=true VERBOSITY=WARNING”.

Filtered BAM files were indexed with samtools index and were used as input in the bamCoverage command of deeptools (v3.0.1)(Ramírez et al., 2014) in order to generate bigwig files. bamCoverage was used with the options – binSize 1 –normalizeUsing RPGC – effectiveGenomeSize 2150570000 –extendReads –outFileFormat bigwig. In order to call peaks, bam files of each timepoint and experiment were merged using the samtools merge command. Resulting merged bam files were indexed, and peaks were called using MACS2 with the options -f BAMPE –nolambda –nomodel -g mm -q 0.05.

### Determination of differentially accessible ATAC peaks

In order to pinpoint regions of interest, peaks of all timepoints and all experiments were merged using the bedtools suite command bedtools merge. Read counts falling within the merged peak regions were calculated using the Rsubread package and the featurecounts command with the options isPairedEnd=T, strandSpecific=0, useMetaFeatures=F. For each transdifferentiation experiment, DESeq2 was used in order to compare all timepoints with timepoint 0h. Any peak showing a logFC ≥ 1 & Adjusted p-value ≤ 0.05 & average counts across timepoints ≥ 5 was termed as a differentially accessible region and was kept for further analyses. The total number of peaks isolated was 91830. Variance stabilized counts were calculated for the isolated regions using the DESeq2 command varianceStabilizingTransformation and the options ‘’blind=T’’, fitType=’’parametric’’. Variance stabilized counts were averaged across timepoint replicates by raising them at the power of 2, extracting their average and log2 transforming the resulting mean. PCA was applied to this dataset using the R prcomp command, with “scale=T”.

To group peaks, PCA was initially applied and PC1 and PC2 values for the 91,830 regions were used in order to arbitrary cluster peaks into 3 groups depending on the sign of their PC1 and PC2 values. Values for each of the 3 groups were visualized using the pheatmap package. Visual examination of the 3 main groups showed different trends: Peaks whose accessibility is higher at 120h (43429 peaks), is lower at 120h (36380 peaks) and is higher at 18h (12021 peaks).

### Motif analysis

Peaks from the 3 different groups were centered and extended 50bp upstream and downstream. Nucleotide sequences for each centered peak were extracted using bedtools getfasta. Sequences were submitted into MEME-ChIP with the following parameters: -dna -seed 49 - meme-nmotifs 20 -meme-minw 5 -meme-minsites 2 -meme-minw 4 -meme-maxw 12. TOMTOM was run using the output meme.txt file in order to identify matches of known transcription factor motifs to the *de novo* discovered motifs. For each TOMTOM output a series of additional filtering steps were undertaken:

1. *De novo* motif sequences need to have <=75% rate for each nucleotide (filtering out repetitive motifs).
2. TOMTOM q-values have to be <=0.01.
3. The matched transcription factor has to be expressed at least at one timepoint.

### Promoter accessibility analysis

Genomic coordinates of mm10 genes were downloaded from the UCSC table browser (RefSeq genes). A single promoter region was assigned to each gene. The region consisted of 1kb upstream and downstream of the transcription start site of the largest transcript of each gene. Counts for each timepoint and each transdifferentiation experiment were assigned to each promoter as described above. DESeq2 was used in order to identify differentially accessible promoters as described above with the following differences regarding the cutoffs used: FoldChange>=1.5 & p-value <=0.05. Variance stabilized counts were extracted for each differentially accessible promoter, a mean value per replicate was extracted and the values were plotted using pheatmap. Promoters were then grouped into 8 clusters. Baloonplots depict all pathways associated with a specific keyword that were found enriched in at least 1 cluster.

For each promoter cluster and each promoter, log2FC changes were extracted by comparing expression levels (DESeq2 normalized counts) of every timepoint with the corresponding timepoint 0h of the experiment.

### Virtual ChIP

C/EBPα and PU.1 binding profiles from ChIP-seq experiments in mouse B cell to macrophage transdifferentiation system were retrieved from earlier work (Van Oevelen et al., 2015). C/EBPα and PU.1 peaks from timepoints 0h, 3h, 12h and 24h were pooled and merged using the bedtools merge command. Each peak was assigned a unique identifier corresponding to the timepoints and experiments the peak was “present”. 6 different groups of peaks were extracted from the pooled file:

1, 2 and 3. Peaks bound by PU.1 at 0h but not at 24h. Group was split further into two sub-groups depending on whether C/EBPα was found to bind at any timepoint.

4, 5 and 6. Peaks bound by C/EBPα at 24h but not at 0h. Group was split further into two sub-groups depending on whether PU.1 was found to bind at any timepoint.

Three different kinds of plots were used to summarize the accessibility dynamics of the six group of peaks in our transdifferentiation system. For each peak the average ATAC-seq bigwig score was calculated using deeptools multiBigwigSummary. Any peak overlapping with mm10 encode blacklisted regions was excluded. Values were averaged across timepoint replicates and visualized in R using the pheatmap package. The same values used for the heatmap peak values were used. Z-transformed values were calculated for every peak.

### Single molecule tracking (SMT)

30,000 NIH 3T3 cells inducible for CEBPAwt-HALO or CEBPAr35a-HALO were seeded in a LabTek-II chambered 8 well plates (Lab-Tek 155049), and induced for 6h or 24h with 1ug/ml doxycycline, with or without prior infection with TETO-FUW-PU.1 lentivirus infection. Right before imaging, cells were treated with 5nM of Janelia Fluor 549 (JF549) HaloTag ligand (a kind gift from Luke Lavis, HHMI) for 15 minutes. Cells were subsequently washed three times in PBS at 37C, and Phenol Red-free High Glucose medium was added to each well. All imaging was carried out under HILO conditions (Tokunaga et al., 2008). For imaging experiments, one frame was acquired with 100ms of exposure time (10 Hz) to measure the intensity of fluorescence of the nuclei, and in SMT)experiments, 5000 frames were acquired with an exposure of 10ms (100 Hz).

Imaging experiments were carried out in Phenol red-free High Glucose Medium (ThermoFisher, Cat#21063029) pyruvate, GlutaMAX, in an imaging chamber heated at 37°C (more details in the Single Molecule Live Cell Imaging section). All live-cell imaging experiments of SMT were carried out in a Nanoimager S from Oxford Nanoimaging Limited (ONI), in a temperature and humidity-controlled chamber, a scientific Complementary metal–oxide– semiconductor (sCMOS) camera with a 2.3 electrons rms read noise at standard scan, a 100X, 1.49 NA oil immersion objective and a 561 nm green laser. Images were acquired with the Nanoimager software. Quantification and Statistical Analysis of SMT was performed as previously described (Lerner et al., 2020). All scripts are publicly available.

### Two Parameter SMT Tracking Analysis

In brief, TIF stacks SMT movies were analyzed using MATLAB-based SLIMfast script (Teves et al., 2016) a modified version of MTT (Sergé et al., 2008), with a Maximal expected Diffusion Coefficient (DMax) of 3 μm2/s-1. The SLIMfast output .txt files were reorganized by the homemade csv_converter.m MATLAB script (available in (Lerner et al., 2020) in .csv format for further analysis. The single molecule tracking .csv files (see previous section) were first classified by the homemade SMT_Motion_Classifier.m MATLAB script. Single molecule trajectories (or tracks) with a track duration shorter than 5 frames were discarded from the analysis. Motion tracks are classified by the script in different groups: tracks with α ≤ 0.7 were considered as Confined; motion tracks with 0.7 < α < 1 as Brownian; and motion tracks with α ≥ 1 as Directed. In addition, the motion tracks showing a behavior similar to a levy-flight (presenting mixed Confined and Directed/Brownian behavior) were detected by the presence of a jump superior to the average jump among the track + a jump threshold of 1.5, and classified as “Butterfly.” Butterfly motion tracks were segmented into their corresponding Confined and Directed/Brownian sub-trajectories for posterior analysis. As an additional filtering step of confined motions (including confined segments of Butterfly tracks), we defined a jump threshold of 100nm, to filter out motion tracks with an average frame-to-frame jump size larger than 100nm.

### Data mining of published datasets

DNA-binding peaks of C/EBPα and PU.1 during BMT were extracted from (Van Oevelen et al. 2015) and analysed as stated above. Single-cell expression trajectories and correlations in B cell transdifferentiation and reprogramming were processed from (Francesconi et al., 2019). Gene expression data from hematopoietic cells (CMP, GMP, Monocyte and Granulocyte (neutrophil)) were from (Ohlsson et al., 2016).

### Statistical analyses

Statistical analyses were performed using Prism 9 software. To calculate significance, samples from at least 3 biologically independent experiments were analyzed. Two biological replicates were used for RNA- and ATAC-sequencing experiments and statistics applied to the expression of a collection of genes. For samples with n ≥ 3, values shown in the figures represent mean ± standard deviation. Box plots represent median with quartiles and whiskers and individual values are shown. One-way, two-way ANOVA (with the corresponding multiple comparison analyses) and Student’s t-tests were applied accordingly. P-values appear indicated in each figure only when ≤ 0.05. In time-course experiments, p-values of differences between conditions by two-way ANOVA are shown. In box plots, p-values of each individual timepoint as well as p-values of differences between conditions by two-way ANOVA are shown.

---

**Table S1.** List of antibodies used for cell sorting and Flow cytometry experiments.

| FACS/Cell sorting |  |  |  |  |
| --- | --- | --- | --- | --- |
| Antibody | Company | Catalogue | Species | Dilution |
| Cd16/Cd32<br>(FcBlock) | BD Pharmingen | 553142 | Rat | 1:400 |
| Cd19-Biotin | BD Biosciences | 553784 | Rat | 1:400 |
| Mac1-Biotin | BD Pharmingen | 557395 | Rat | 1:400 |
| hCD4-Biotin | eBioscience | 13-0049 | Mouse | 1:33 |
| Cd19-APC | BD Pharmingen | 550992 | Rat | 1:400 |
| Mac1-PE-Cy7 | BD Pharmingen | 552850 | Rat | 1:400 |
| Ly6g-PE | Pharmingen | 553128 | Rat | 1:400 |
| Mac1-APC | eBioscience | 17-0112-83 | Rat | 1:400 |
| hCD4-PE | BD Pharmingen | 555347 | Mouse | 1:20 |
| hCD16/CD32<br>(hFcBlock) | Invitrogen | 16-9161-73 | - | 1:20 |
| hCD19-APC-Cy7 | BD Pharmingen | 557791 | Mouse | 1:33 |
| hMac1-APC | BD Pharmingen | 561015 | Mouse | 1:33 |
| Cd16/Cd32-FITC | BD Pharmingen | 553144 | Rat | 1:400 |
| cKit-APC-Cy7 | Invitrogen | 47-1172-82 | Rat | 1:400 |
| Cd34-APC | BD Pharmingen | 560230 | Rat | 1:50 |
| Sca1-PE-Cy7 | BD Pharmingen | 558162 | Rat | 1:400 |
| Sca1-PerCP-Cy5.5 | eBioscience | 35-5981-82 | Rat | 1:400 |
| Cd41-PE-Cy7 | eBioscience | 25-0411-82 | Rat | 1:400 |

**Table S2.**
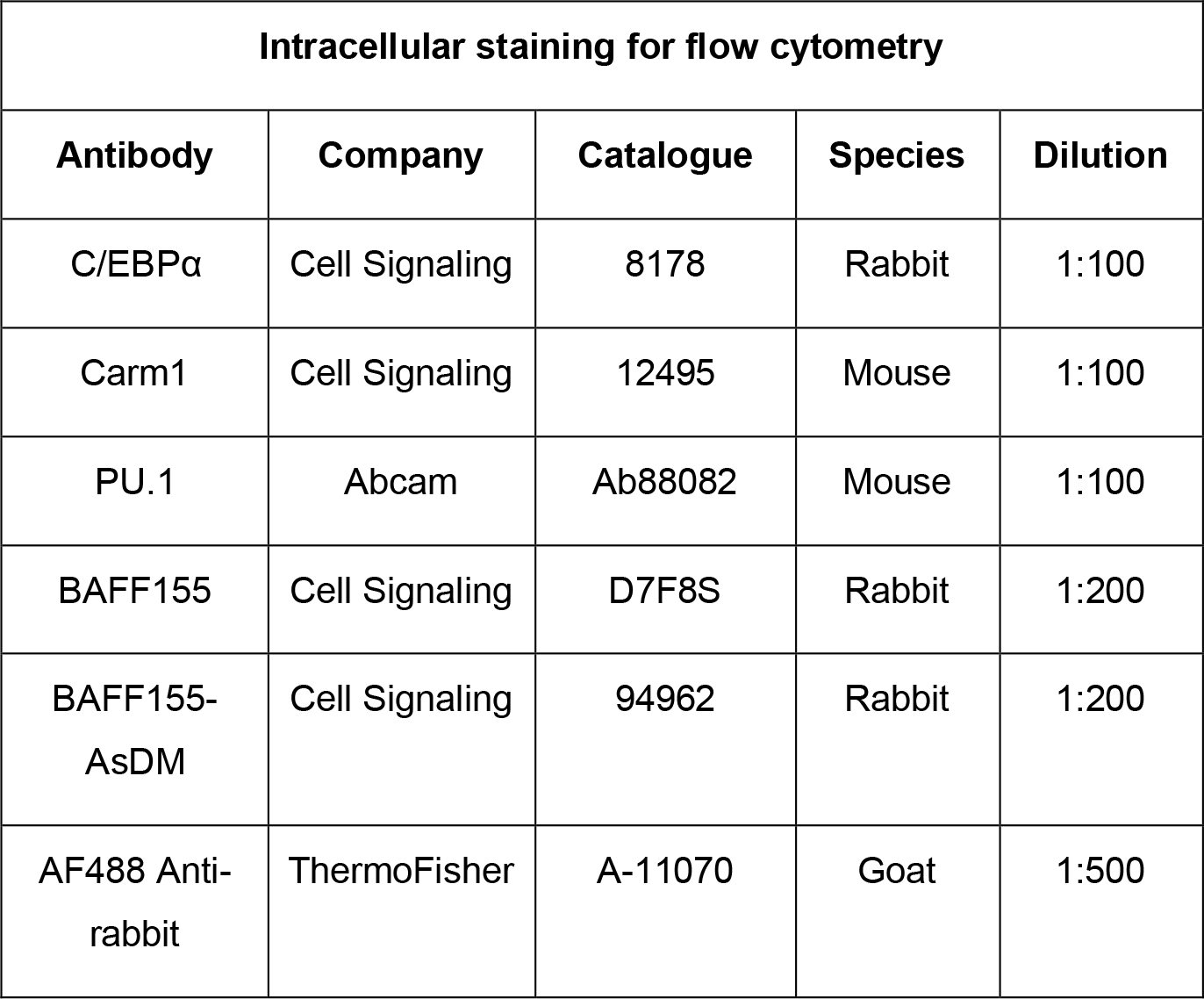
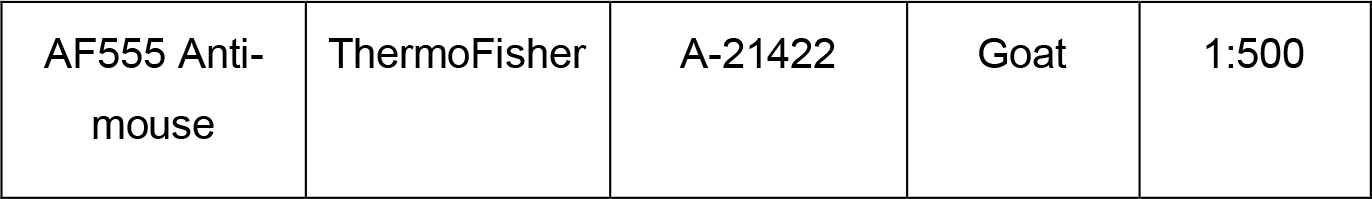
List of antibodies and fluorochromes used for immunofluorescence and intracellular staining for flow cytometry.

| Intracellular staining for flow cytometry |  |  |  |  |
| --- | --- | --- | --- | --- |
| Antibody | Company | Catalogue | Species | Dilution |
| C/EBPα | Cell Signaling | 8178 | Rabbit | 1:100 |
| Carm1 | Cell Signaling | 12495 | Mouse | 1:100 |
| PU.1 | Abcam | Ab88082 | Mouse | 1:100 |
| BAFF155 | Cell Signaling | D7F8S | Rabbit | 1:200 |
| BAFF155-AsDM | Cell Signaling | 94962 | Rabbit | 1:200 |
| AF488 Anti-<br>rabbit | ThermoFisher | A-11070 | Goat | 1:500 |
| AF555 Anti-mouse | ThermoFisher | A-21422 | Goat | 1:500 |

**Table S3.** Chemical reagents used to prepare buffers for western blot.

| Running buffer | Transfer buffer | TBST |
| --- | --- | --- |
| 25mM Tris-base | 25mM Tris-HCl<br>pH=3.8 | 10mM Tris HCl=7.5 |
| 200mM glycine | 200mM glycine | 100mM NaCl |
| 0.1% SDS | 20% methanol | 0.1% Tween 20 |

**Table S4.** List of antibodies used for western blot experiments.

| Antibody | Company | Catalogue | Species | Dilution |
| --- | --- | --- | --- | --- |
| C/EBPα | Cell Signaling | 8178 | Rabbit | 1:1000 |
| aDMA | Cell Signaling | 13522S | Rabbit | 1:1000 |
| aDMA | Upstate | #07-414 | Rabbit | 1:1000 |
| HA | Covance | #MMS-101R | Mouse | 1:1000 |
| Flag | Sigma | F3165 | Mouse | 1:1000 |
| Flag | Abnova | PAB 29056 | Chicken | 1:1000 |
| BAFF155 | Cell Signaling | D7F8S | Rabbit | 1:1000 |
| BAFF155-AsDM | Cell Signaling | 94962 | Rabbit | 1:1000 |
| PU.1 | Abcam | Ab88082 | Mouse | 1:1000 |
| Vinculin | Merck | V9131 | Mouse | 1:200 |
| Gapdh | Abcam | Ab8245 | Mouse | 1:5000 |
| H3 | Abcam | Ab10799 | Mouse | 1:1000 |

**Table S5.**
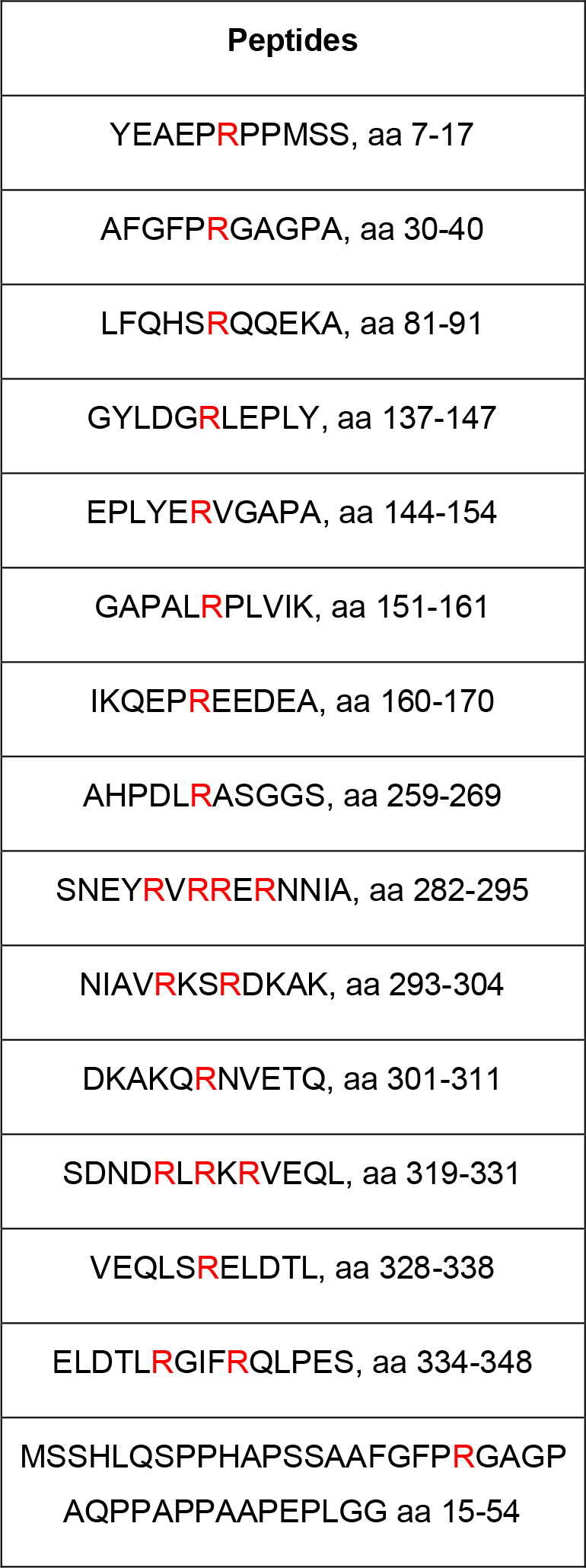
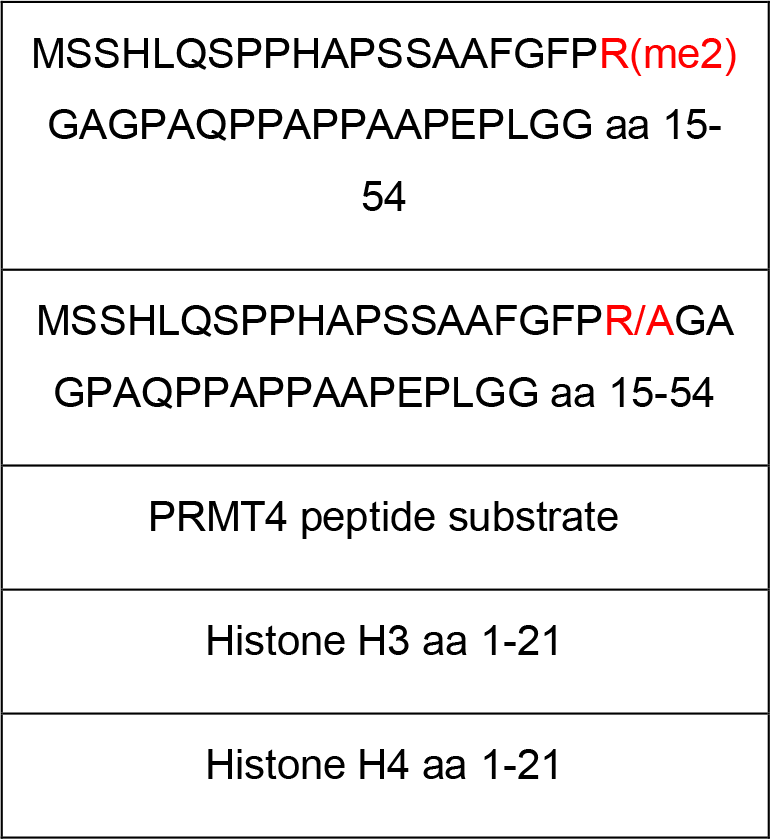
List of peptides used for in vitro methylation experiments.

**Table S6.**
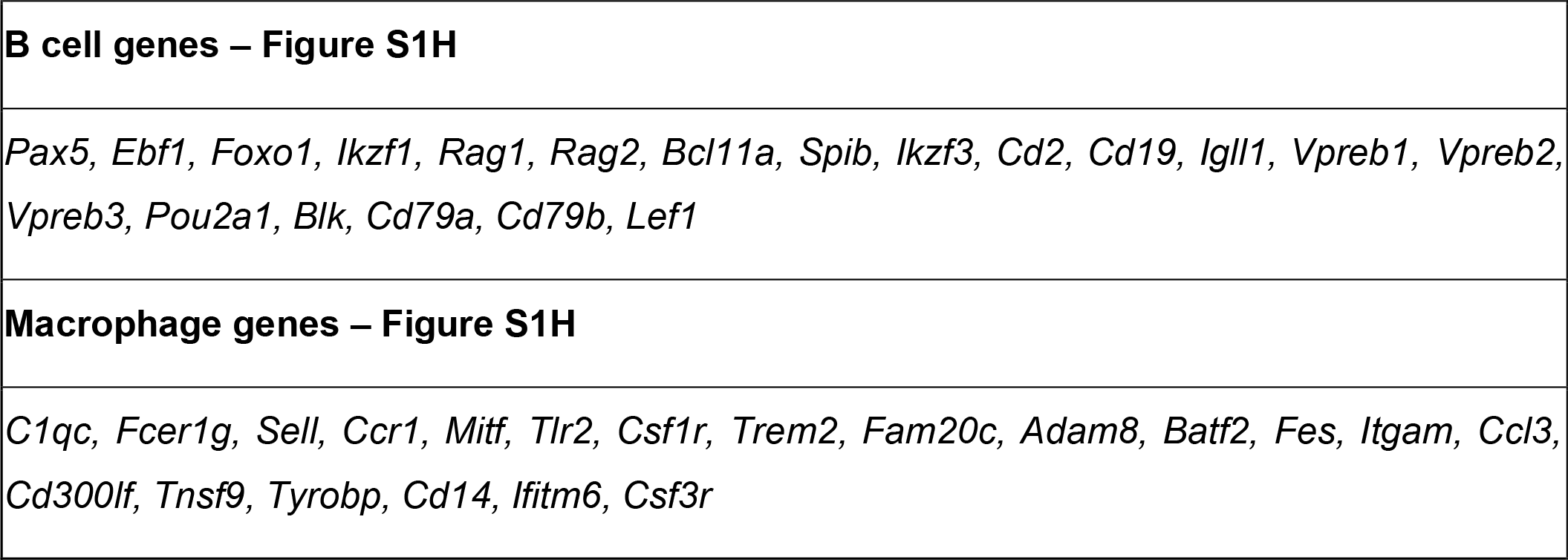
List of genes used to analyze kinetics of specific signatures.

## Newly Created Materials

The new constructs and cell lines listed can be requested from the corresponding authors. The sequencing data will be deposited at GEO and made freely available

## REFERENCES

Arinobu Y, Mizuno S, Chong Y, Shigematsu H, Iino T, Iwasaki H, Graf T, Mayfield R, Chan S, Kastner P, Akashi K. 2007. Reciprocal Activation of GATA-1 and PU.1 Marks Initial Specification of Hematopoietic Stem Cells into Myeloerythroid and Myelolymphoid Lineages. Cell Stem Cell 1:416–427. doi:10.1016/j.stem.2007.07.004

Bedford MT, Clarke SG. 2009. Protein Arginine Methylation in Mammals: Who, What, and Why. Mol Cell 33:1–13. doi:10.1016/j.molcel.2008.12.013

Bedford MT, Richard S. 2005. Arginine methylation: An emerging regulator of protein function. Mol Cell 18:263–272. doi:10.1016/j.molcel.2005.04.003

Buenrostro JD, Wu B, Chang HY, Greenleaf WJ. 2015. ATAC-seq: A method for assaying chromatin accessibility genome-wide. Curr Protoc Mol Biol 2015:21.29.1–21.29.9. doi:10.1002/0471142727.mb2129s109

Bussmann LH, Schubert A, Vu Manh TP, De Andres L, Desbordes SC, Parra M, Zimmermann T, Rapino F, Rodriguez-Ubreva J, Ballestar E, Graf T. 2009. A Robust and Highly Efficient Immune Cell Reprogramming System. Cell Stem Cell 5:554–566. doi:10.1016/j.stem.2009.10.004

Cai DH, Wang D, Keefer J, Yeamans C, Hensley K, Friedman AD. 2008. C/EBPα:AP-1 leucine zipper heterodimers bind novel DNA elements, activate the PU.1 promoter and direct monocyte lineage commitment more potently than C/EBPα homodimers or AP-1. Oncogene 27:2772–2779. doi:10.1038/SJ.ONC.1210940

Chang NC, Sincennes MC, Chevalier FP, Brun CE, Lacaria M, Segalés J, Muñoz-Cánoves P, Ming H, Rudnicki MA. 2018. The Dystrophin Glycoprotein Complex Regulates the Epigenetic Activation of Muscle Stem Cell Commitment. Cell Stem Cell 22:755–768.e6. doi:10.1016/j.stem.2018.03.022

Chen J, Zhang Z, Li L, Chen BC, Revyakin A, Hajj B, Legant W, Dahan M, Lionnet T, Betzig E, Tjian R, Liu Z. 2014. Single-molecule dynamics of enhanceosome assembly in embryonic stem cells. Cell 156:1274–1285. doi:10.1016/j.cell.2014.01.062

Choi J, Baldwin TM, Wong M, Bolden JE, Fairfax KA, Lucas EC, Cole R, Biben C, Morgan C, Ramsay KA, Ng AP, Kauppi M, Corcoran LM, Shi W, Wilson N, Wilson MJ, Alexander WS, Hilton DJ, De Graaf CA. 2019. Haemopedia RNA-seq: A database of gene expression during haematopoiesis in mice and humans. Nucleic Acids Res 47:D780–D785. doi:10.1093/nar/gky1020

Deribe YL, Pawson T, Dikic I. 2010. Post-translational modifications in signal integration. Nat Struct Mol Biol 17:666–672. doi:10.1038/nsmb.1842

Di Stefano B, Sardina JL, Van Oevelen C, Collombet S, Kallin EM, Vicent GP, Lu J, Thieffry D, Beato M, Graf T. 2014. C/EBPα poises B cells for rapid reprogramming into induced pluripotent stem cells. Nature 506:235–239. doi:10.1038/nature12885

Dobin A, Davis CA, Schlesinger F, Drenkow J, Zaleski C, Jha S, Batut P, Chaisson M, Gingeras TR. 2013. STAR: Ultrafast universal RNA-seq aligner. Bioinformatics 29:15–21. doi:10.1093/bioinformatics/bts635

Ebisuya M, Briscoe J. 2018. What does time mean in development? Development 145. doi:10.1242/dev.164368

Eyquem S, Chemin K, Fasseu M, Chopin M, Sigaux F, Cumano A, Bories JC. 2004. The development of early and mature B cells is impaired in mice deficient for the Ets-1 transcription factor. Eur J Immunol 34:3187–3196. doi:10.1002/EJI.200425352

Feng R, Desbordes SC, Xie H, Tillo ES, Pixley F, Stanley ER, Graf T. 2008. PU.1 and C/EBPα/β convert fibroblasts into macrophage-like cells. Proceedings of the National Academy of Sciences 105:6057–6062. doi:10.1073/pnas.0711961105

Fernandez Garcia M, Moore CD, Schulz KN, Alberto O, Donague G, Harrison MM, Zhu H, Zaret KS. 2019. Structural Features of Transcription Factors Associating with Nucleosome Binding. Mol Cell 75:921–932.e6. doi:10.1016/j.molcel.2019.06.009

Francesconi M, di Stefano B, Berenguer C, Andrés-Aguayo L de, Plana-Carmona M, Mendez-Lago M, Guillaumet-Adkins A, Rodriguez-Esteban G, Gut M, Gut IG, Heyn H, Lehner B, Graf T. 2019. Single cell RNA-seq identifies the origins of heterogeneity in efficient cell transdifferentiation and reprogramming. Elife 8:1–22. doi:10.7554/eLife.41627

Graf T, Enver T. 2009. Forcing cells to change lineages. Nature 462:587–594. doi:10.1038/nature08533

Greenblatt SM, Man N, Hamard PJ, Asai T, Karl D, Martinez C, Bilbao D, Stathais V, McGrew-Jermacowicz A, Duffort S, Tadi M, Blumenthal E, Newman S, Vu L, Xu Y, Liu F, Schurer SC, McCabe MT, Kruger RG, Xu M, Yang FC, Tenen D, Watts J, Vega F, Nimer SD. 2018. CARM1 Is Essential for Myeloid Leukemogenesis but Dispensable for Normal Hematopoiesis. Cancer Cell 33:1111–1127.e5. doi:10.1016/j.ccell.2018.05.007

Guo Z, Zheng L, Xu H, Dai H, Zhou M, Pascua MR, Chen QM, Shen B. 2010. Methylation of FEN1 suppresses nearby phosphorylation and facilitates PCNA binding. Nat Chem Biol 6:766–773. doi:10.1038/NCHEMBIO.422

Heath V, Suh HC, Holman M, Renn K, Gooya JM, Parkin S, Klarmann KD, Ortiz M, Johnson P, Keller J. 2004. C/EBPα deficiency results in hyperproliferation of hematopoietic progenitor cells and disrupts macrophage development in vitro and in vivo. Blood 104:1639–1647. doi:10.1182/blood-2003-11-3963

Heinz S, Benner C, Spann N, Bertolino E, Lin YC, Laslo P, Cheng JX, Murre C, Singh H, Glass CK. 2010. Simple Combinations of Lineage-Determining Transcription Factors Prime cis-Regulatory Elements Required for Macrophage and B Cell Identities. Mol Cell 38:576–589. doi:10.1016/j.molcel.2010.05.004

Hertweck A, Mucha MV De, Barber PR, Dagil R, Porter H, Ramos A, Lord GM, Jenner RG. 2022. The TH1 cell lineage-determining transcription factor T-bet suppresses TH2 gene expression by redistributing GATA3 away from TH2 genes. Nucleic Acids Res 1. doi:https://doi.org/10.1093/nar/gkac258

Hosokawa H, Ungerbäck J, Wang X, Matsumoto M, Nakayama KI, Cohen SM, Tanaka T, Rothenberg E V. 2018. Transcription Factor PU.1 Represses and Activates Gene Expression in Early T Cells by Redirecting Partner Transcription Factor Binding. Immunity 48:1119-1134.e7. doi:10.1016/j.immuni.2018.04.024

Hsiao K, Zegzouti H, Goueli SA. 2016. Methyltransferase-Glo: A universal, bioluminescent and homogenous assay for monitoring all classes of methyltransferases. Epigenomics 8:321–339. doi:10.2217/EPI.15.113

Hu C-J, Rao S, Ramirez-Bergeron DL, Garrett-Sinha LA, Gerondakis S, Clark MR, Simo MC. 2001. PU.1/Spi-B Regulation of c-rel Is Essential for Mature B Cell Survival. Immunity 15:545–555.

Jack I, Seshadri R, Garson M, Michael P, Callen D, Zola H, Morley A. 1986. RCH-ACV: A lymphoblastic leukemia cell line with chromosome translocation 1;19 and trisomy 8. Cancer Genet Cytogenet 19:261–269. doi:10.1016/0165-4608(86)90055-5

Kawabe YI, Wang YX, McKinnell IW, Bedford MT, Rudnicki MA. 2012. Carm1 regulates Pax7 transcriptional activity through MLL1/2 recruitment during asymmetric satellite stem cell divisions. Cell Stem Cell 11:333–345. doi:10.1016/j.stem.2012.07.001

Kim D, Lee J, Cheng D, Li J, Carter C, Richie E, Bedford MT. 2010. Enzymatic activity is required for the in Vivo functions of CARM1. Journal of Biological Chemistry 285:1147–1152. doi:10.1074/jbc.M109.035865

Klemm SL, Shipony Z, Greenleaf WJ. 2019. Chromatin accessibility and the regulatory epigenome. Nature Reviews 20:207–220. doi:10.1038/s41576-018-0089-8

Konstantinides N, Holguera I, Rossi AM, Escobar A, Dudragne L, Chen Y-C, Tran TN, Martínez Jaimes AM, Özel MN, Simon F, Shao Z, Tsankova NM, Fullard JF, Walldorf U, Roussos P, Desplan C. 2022. A complete temporal transcription factor series in the fly visual system. Nature 604:316–322. doi:10.1038/s41586-022-04564-w

Kowenz-Leutz E, Pless O, Dittmar G, Knoblich M, Leutz A. 2010. Crosstalk between C/EBPΒ phosphorylation, arginine methylation, and SWI/SNF/Mediator implies an indexing transcription factor code. EMBO Journal 29:1105–1115. doi:10.1038/emboj.2010.3

Kowenz-Leutz E, Twamley G, Ansieau S, Leutz A. 1994. Novel mechanism of C/EBPβ (NF-M) transcriptional control: Activation through derepression. Genes Dev 8:2781–2791. doi:10.1101/gad.8.22.2781

Kueh HY, Champhekar A, Nutt SL, Elowitz MB, Rothenberg E v. 2013. Positive Feedback Between PU.1 and the Cell Cycle Controls Myeloid Differentiation. Science (1979) 341:670– 673. doi:10.1126/science.1240831

Laiosa C V., Stadtfeld M, Xie H, de Andres-Aguayo L, Graf T. 2006. Reprogramming of Committed T Cell Progenitors to Macrophages and Dendritic Cells by C/EBPα and PU.1 Transcription Factors. Immunity 25:731–744. doi:10.1016/j.immuni.2006.09.011

Leddin M, Perrod C, Hoogenkamp M, Ghani S, Assi S, Heinz S, Wilson NK, Follows G, Schönheit J, Vockentanz L, Mosammam AM, Chen W, Tenen DG, Westhead DR, Göttgens B, Bonifer C, Rosenbauer F. 2011. Two distinct auto-regulatory loops operate at the PU.1 locus in B cells and myeloid cells. Blood 117:2827–2838. doi:10.1182/blood-2010-08-302976

Lerner J, Gomez-Garcia PA, McCarthy RL, Liu Z, Lakadamyali M, Zaret KS. 2020. Two-Parameter Mobility Assessments Discriminate Diverse Regulatory Factor Behaviors in Chromatin. Mol Cell 79:677–688. doi:10.1016/j.molcel.2020.05.036

Li H, Handsaker B, Wysoker A, Fennell T, Ruan J, Homer N, Marth G, Abecasis G, Durbin R. 2009. The Sequence Alignment/Map format and SAMtools. Bioinformatics 25:2078–2079. doi:10.1093/bioinformatics/btp352

Li J, Zhao Z, Carter C, Ehrlich LIR, Bedford MT, Richie ER. 2013. Coactivator-Associated Arginine Methyltransferase 1 Regulates Fetal Hematopoiesis and Thymocyte Development. The Journal of Immunology 190:597–604. doi:10.4049/jimmunol.1102513

Liu Z, Tjian R. 2018. Visualizing transcription factor dynamics. Journal of Cell Biology 217:1181– 1191. doi:10.1083/jcb.201710038

Love MI, Huber W, Anders S. 2014. Moderated estimation of fold change and dispersion for RNA-seq data with DESeq2. Genome Biol 15:1–21. doi:10.1186/s13059-014-0550-8

Ma O, Hong S, Guo H, Ghiaur G, Friedman AD. 2014. Granulopoiesis Requires Increased C/EBPα Compared to Monopoiesis, Correlated with Elevated Cebpa in Immature G-CSF Receptor versus M-CSF Receptor Expressing Cells. PLoS One 9:e95784. doi:10.1371/journal.pone.0095784

Martin M. 2011. Cutadapt removes adapter sequences from high-throughput sequencing reads. EMBnet J 17.1

Moris N, Pina C, Arias AM. 2016. Transition states and cell fate decisions in epigenetic landscapes. Nat Rev Genet 17:693–703. doi:10.1038/nrg.2016.98

Nakayama K, Szewczyk MM, dela Sena C, Wu H, Dong A, Zeng H, Li F, Ferreira De Freitas R, Eram MS, Schapira M, Baba Y, Kunitomo M, Cary DR, Tawada M, Ohashi A, Imaeda Y, Singh Saikatendu K, Grimshaw CE, Vedadi M, Arrowsmith CH, Barsyte-Lovejoy D, Kiba A, Tomita D, Brown PJ. 2018. TP-064, a potent and selective small molecule inhibitor of PRMT4 for multiple myeloma. Oncotarget 9:18480–18493.

Nerlov C. 2007. The C/EBP family of transcription factors: a paradigm for interaction between gene expression and proliferation control. Trends Cell Biol 17:318–324. doi:10.1016/j.tcb.2007.07.004

Notta F, Zandi S, Takayama N, Dobson S, Gan OI, Wilson G, Kaufmann KB, McLeod J, Laurenti E, Dunant CF, McPherson JD, Stein LD, Dror Y, Dick JE. 2016. Distinct routes of lineage development reshape the human blood hierarchy across ontogeny. Science (1979) 351. doi:10.1126/science.aab2116

Ohlsson E, Schuster MB, Hasemann M, Porse BT. 2016. The multifaceted functions of C/EBPα in normal and malignant haematopoiesis. Leukemia. doi:10.1038/leu.2015.324

Okawa S, Saltó C, Ravichandran S, Yang S, Toledo EM, Arenas E, Del Sol A. 2018. Transcriptional synergy as an emergent property defining cell subpopulation identity enables population shift. Nat Commun 9:1–10. doi:10.1038/s41467-018-05016-8

Orkin SH, Zon LI. 2008. Hematopoiesis: An Evolving Paradigm for Stem Cell Biology. Cell 132:631–644. doi:10.1016/j.cell.2008.01.025

Ramberger E, Sapozhnikova V, Kowenz-Leutz E, Zimmermann K, Nicot N, Nazarov P v., Perez-Hernandez D, Reimer U, Mertins P, Dittmar G, Leutz A. 2021. PRISMA and BioID disclose a motifs-based interactome of the intrinsically disordered transcription factor C/EBPα. iScience 24:102686. doi:10.1016/j.isci.2021.102686

Ramírez F, Dündar F, Diehl S, Grüning BA, Manke T. 2014. DeepTools: A flexible platform for exploring deep-sequencing data. Nucleic Acids Res 42:187–191. doi:10.1093/nar/gku365

Rapino F, Robles EF, Richter-Larrea JA, Kallin EM, Martinez-Climent JA, Graf T. 2013. C/EBPα Induces Highly Efficient Macrophage Transdifferentiation of B Lymphoma and Leukemia Cell Lines and Impairs Their Tumorigenicity. Cell Rep 3:1153–1163. doi:10.1016/j.celrep.2013.03.003

Raschke WC, Baird S, Ralph P, Nakoinz I. 1978. Functional macrophage cell lines transformed by abelson leukemia virus. Cell 15:261–267. doi:10.1016/0092-8674(78)90101-0

Raudvere U, Kolberg L, Kuzmin I, Arak T, Adler P, Peterson H, Vilo J. 2019. G:Profiler: A web server for functional enrichment analysis and conversions of gene lists (2019 update). Nucleic Acids Res 47:W191–W198. doi:10.1093/nar/gkz369

Rayon T, Stamataki D, Perez-Carrasco R, Garcia-Perez L, Barrington C, Melchionda M, Exelby K, Lazaro J, Tybulewicz VLJ, Fisher EMC, Briscoe J. 2020. Species-specific pace of development is associated with differences in protein stability. Science (1979) 369. doi:10.1126/science.aba7667

Schreiber E, Matthias P, Müller MM, Schaffner W. 1989. Rapid detection of octamer binding proteins with ‘mini extracts’, prepared from a small number of cells. Nucleic Acids Res 17:6419. doi:10.1093/nar/17.15.6419

Scott EW, Simon MC, Anastasi J, Singh H. 1994. Requirement of transcription factor PU.1 in the development of multiple hematopoietic lineages. Science (1979) 265:1573–1577. doi:10.1126/science.8079170

Sergé A, Bertaux N, Rigneault H, Marguet D. 2008. Dynamic multiple-target tracing to probe spatiotemporal cartography of cell membranes. Nature Methods 2008 5 :8 5:687–694. doi:10.1038/nmeth.1233

Singh H, Dekoter RP, Walsh JC. 1999. PU.1, a Shared Transcriptional Regulator of Lymphoid and Myeloid Cell Fates. Cold Spring Harb Symp Quant Biol 64:13–20. doi:10.1101/sqb.1999.64.13

Springer T, Galfré G, Secher DS, Milstein C. 1979. Mac-1: a macrophage differentiation antigen identified by monoclonal antibody. Eur J Immunol 9:301–306. doi:10.1002/eji.1830090410

Stoilova B, Kowenz-Leutz E, Scheller M, Leutz A. 2013. Lymphoid to Myeloid Cell Trans-Differentiation Is Determined by C/EBPβ Structure and Post-Translational Modifications. PLoS One 8:e65169. doi:10.1371/journal.pone.0065169

Suresh S, Huard S, Dubois T. 2021. CARM1/PRMT4: Making Its Mark beyond Its Function as a Transcriptional Coactivator. Trends Cell Biol 31:402–417. doi:10.1016/j.tcb.2020.12.010

Teves SS, An L, Hansen AS, Xie L, Darzacq X, Tjian R. 2016. A dynamic mode of mitotic bookmarking by transcription factors. Elife 5. doi:10.7554/ELIFE.22280

Torcal Garcia G, Graf T. 2021. The transcription factor code: a beacon for histone methyltransferase docking. Trends Cell Biol 31:792–800. doi:10.1016/j.tcb.2021.04.001

Torres-Padilla ME, Parfitt DE, Kouzarides T, Zernicka-Goetz M. 2007. Histone arginine methylation regulates pluripotency in the early mouse embryo. Nature 445:214–218. doi:10.1038/nature05458

van Oevelen C, Collombet S, Vicent G, Hoogenkamp M, Lepoivre C, Badeaux A, Bussmann L, Sardina JL, Thieffry D, Beato M, Shi Y, Bonifer C, Graf T. 2015. C/EBPα Activates Pre-existing and de Novo Macrophage Enhancers during Induced Pre-B Cell Transdifferentiation and Myelopoiesis. Stem Cell Reports 5:232–247. doi:10.1016/j.stemcr.2015.06.007

Velten L, Haas SF, Raffel S, Blaszkiewicz S, Hennig BP, Hirche C, Lutz C, Buss EC, Boch T, Hofmann W, Ho AD, Huber W. 2017. Human haematopoietic stem cell lineage commitment is a continuous process 19:271–281. doi:10.1038/ncb3493.Human

Wang K, Wei G, Liu D. 2012. CD19: a biomarker for B cell development, lymphoma diagnosis and therapy. Exp Hematol Oncol 1:36. doi:10.1186/2162-3619-1-36

Wang L, Zhao Z, Meyer MB, Saha S, Yu M, Guo A, Wisinski KB, Huang W, Cai W, Pike JW, Yuan M, Ahlquist P, Xu W. 2014. CARM1 methylates chromatin remodeling factor BAF155 to enhance tumor progression and metastasis. Cancer Cell 25:21–36. doi:10.1016/j.ccr.2013.12.007

Wu Q, Schapira M, Arrowsmith CH, Barsyte-Lovejoy D. 2021. Protein arginine methylation: from enigmatic functions to therapeutic targeting. Nat Rev Drug Discov 20:509–530. doi:10.1038/s41573-021-00159-8

Xie H, Ye M, Feng R, Graf T. 2004. Stepwise reprogramming of B cells into macrophages. Cell 117:663–676. doi:10.1016/S0092-8674(04)00419-2

Xue HH, Bollenbacher-Reilley J, Wu Z, Spolski R, Jing X, Zhang YC, McCoy JP, Leonard WJ. 2007. The Transcription Factor GABP Is a Critical Regulator of B Lymphocyte Development. Immunity 26:421–431. doi:10.1016/J.IMMUNI.2007.03.010

Yadav N, Cheng D, Richard S, Morel M, Iyer VR, Aldaz CM, Bedford MT. 2008. CARM1 promotes adipocyte differentiation by coactivating PPARγ. EMBO Rep 9:193–198. doi:10.1038/sj.embor.7401151

Zhang P, Iwasaki-Arai J, Iwasaki H, Fenyus ML, Dayaram T, Owens BM, Shigematsu H, Levantini E, Huettner CS, Lekstrom-Himes JA, Akashi K, Tenen DG. 2004. Enhancement of hematopoietic stem cell repopulating capacity and self-renewal in the absence of the transcription factor C/EBPα. Immunity 21:853–863. doi:10.1016/j.immuni.2004.11.006

Zhang XK, Moussa O, LaRue A, Bradshaw S, Molano I, Spyropoulos DD, Gilkeson GS, Watson DK. 2008. The Transcription Factor Fli-1 Modulates Marginal Zone and Follicular B Cell Development in Mice. The Journal of Immunology 181:1644–1654. doi:10.4049/JIMMUNOL.181.3.1644

